# Adaptive evolution of engineered *Saccharomyces cerevisiae* in favored and unusual chemical environments

**DOI:** 10.1101/2025.10.29.685190

**Authors:** Natalia Kakko von Koch, Olli Lohilahti, Andreas Møllerhøj Vestergaard, An Nguyen, Tomas Strucko, Paula Jouhten

## Abstract

Engineered microbial cells can produce a wide range of industrially relevant chemicals such as pharmaceuticals, fuels, and material precursors. The use of microbial cells for chemical production from renewable resources could replace oil-based chemistry and contribute to tackling global grand challenges of climate warming and resource insufficiency. However, it is underexplored how the chemical production by engineered microbial cells is affected by them being proliferating catalysts exposed to Darwinian selection. All proliferating cells are unavoidably subjected to Darwinian selection which favors fitness beneficial phenotypes that seldom include engineered chemical production.

Here, adaptive laboratory evolution was performed to characterize the effect of Darwinian selection on *Saccharomyces cerevisiae* strains expressing two different heterologous pigment producing pathways, blue-coloured indigoidine and red-coloured bikaverin. *S. cerevisiae* haploid S288C based strain had the genes for bikaverin synthesis integrated in the same locus as the genes for indigoidine synthesis in haploid and diploid *S. cerevisiae* CEN.PK-based strains. The two different pigment producing strains were cultivated in rich and synthetic defined (without amino acids) media with respirative galactose as the sole carbon source for ∼200 and ∼175 generations, respectively. While CEN.PK-based lineages rapidly lost indigoidine pigmentation independent of growth medium or ploidy, bikaverin pigmentation in S288C-based lineages was robust. The adaptive solutions detected in S288C-based bikaverin producing lineages involved mutations in the galactose utilization pathway whereas the heterologous indigoidine pathway was recurrently mutated in the corresponding lineages. When the bikaverin producing S288C-based lineages were adaptively evolved on the favored glucose carbon source instead, pigmentation declined. Thus, the robustness of the engineered traits appears dependent on challenges in production environment and availability and fitness benefits of adaptive solutions.

Whether or when engineered traits of microbial cells are robust when they proliferate in industrial use has scarcely been assessed. Here light was shed to the factors affecting the adaptive loss of engineered traits to facilitate the development of strains and biotechnological processes, including chemical environments, for robust long-term production.

## Introduction

Engineered microbial cells can produce a wide range of industrially relevant chemicals such as pharmaceuticals, fuels, and material precursors (Kim et al., 2023). Oil-based chemistry could be replaced by using microbial cells for chemical production from renewable resources and this would contribute to tackling global grand challenges of climate warming and resource insufficiency. However, the chemical production using microbial cells may not be stable which has a negative influence on the feasibility of biotechnological processes. Microbial cells as chemical producers are living catalysts that are influenced by Darwinian selection. Darwinian selection acts on all proliferating cultures of cells selecting for fitness beneficial phenotypes. Cells engineered for the production of heterologous chemical seldom exhibit fitness beneficial phenotypes. Even when the target compound is not toxic, engineered production competes for resources such as translational machinery, precursors, energy and redox power with cell growth (Kim et al., 2013; Scott et al., 2014; Wu et al., 2016; Zhang et al., 2022). This causes burden which is usually observed as reduced growth rate and yield.

Producer cells grow not only during the production process but also in strain maintenance and biomass expansion for industrial scale processes. Random mutations occur during growth-associated DNA replication at a rate that may not limit the occurrence of mutations that enable improved growth (particularly if they are not rare) (Arjan G et al., 1999; Hall et al., 2010). Mutational target affecting the production is formed of the heterologous pathway integrated in the genome and native metabolic genes and regulators essential for the provision of precursors, energy and redox power to the production. When a loss of production due to a random mutation in a clone allows improved growth, Darwinian selection will lead to an enrichment of such clones (Csörgő et al., 2012; Munkler et al., 2024; Xie et al., 2018). They also deplete resources from the producing sub-population. As a consequence of the reducing producing sub-population and the resource limitation, the production levels are reduced (Rugbjerg et al., 2018a; Rugbjerg and Sommer, 2019).

Production levels can be reduced also due to phenotypic heterogeneity in response to environmental variation in the growth conditions (i.e., pH, pressure and oxygen levels) or stochastic noise in biological processes, like gene expression (Binder et al., 2017; Gasperotti et al., 2020; Xiao et al., 2016). However, environmental variation can be reduced by process optimization and stochastic noise involves commonly very low and non-increasing proportions of populations (van Heerden et al., 2014; Xiao et al., 2016). In contrast, genetic changes are heritable to daughter cells and therefore higher fitness (i.e., low-producers) clones are unavoidably enriched through Darwinian selection. Thus, production loss due to adaptive evolution forms a more severe risk for biotechnological production processes (Tan et al., 2024). Yet, the evolutionary robustness of engineered, heterologous, pathways have not been sufficiently assessed.

Here, the evolutionary adaptation of engineered yeast *Saccharomyces cerevisiae* strains expressing two different heterologous pigment producing pathways, blue indigoidine and red bikaverin was characterized. Indigoidine competes with protein synthesis on the precursor L-glutamine, whereas the precursor of bikaverin is acetyl-CoA that cells use mainly for lipid biosynthesis. The heterologous genes required for the synthesis of indigoidine and bikaverin were introduced into the same genomic locus in haploid CEN.PK- and S288C-based strains, respectively. Both strains were adaptively evolved in rich and synthetic defined (without amino acids) media with galactose as the sole carbon source. The poor galactose utilizer S288C strain (Mortimer and Johnston, 1986; Otero et al., 2010) was adaptively evolved also on glucose as the sole carbon source. Additionally, a diploid indigoidine producing CEN.PK-based strain was included in the adaptive evolution experiments to assess the effect of two chromosome copies on the adapted genotypes.

## Results

### Growth of S288C-based lineages was notably improved during ALE

To assess engineered trait stability under Darwinian selection, ALE of engineered, heterologous pigment producing, strains was performed (Figure 1). The pathway for blue pigment indigoidine production (Wehrs et al., 2018) was integrated in a haploid CEN.PK113-7D strain (IND-H) and diploid CEN.PK113-1Ax7D strain (IND-D), at well-characterized genomic locus XII-5 (Jessop-Fabre et al., 2016; Mikkelsen et al., 2012). The pathway consisted of two heterologous genes encoding indigoidine synthetase (BpsA) and a 4⍰-phosphopantetheinyl transferase (Sfp). BpsA is a non-ribosomal peptide synthetase (NRPS), and Sfp is required to activate the BpsA from its apoform to its holo-form (Wehrs et al., 2018). After activation, the NRPS condenses two L-glutamine molecules to form a single indigoidine molecule. The heterologous pathway for red pigment bikaverin synthesis was also integrated into the XII-5 locus (Jessop-Fabre et al., 2016; Mikkelsen et al., 2012) but in haploid S288C-based strain (BIK). The pathway consisted of three genes: Bik1, npgA, and Bik2-Bik3 encoding a fusion enzyme found to substantially improve bikaverin production (Zhao et al., 2020). Bik1 encodes a polyketide synthase (PKS) that, when activated with the npgA encoded 4⍰-phosphopantetheinyl transferase, catalyzes the formation of the polyketide backbone of bikaverin, pre-bikaverin. Bik2 encodes for a FAD-dependent monooxygenase that oxidizes pre-bikaverin to oxo-pre-bikaverin. Bik3 encodes for a methyltransferase that catalyzes the methylation of oxo-pre-bikaverin to dehydroxybikaverin. Dehydroxy-bikaverin is then oxidized to bikaverin by Bik2. Alternatively, Bik2 can oxidize the intermediate to nor-bikaverin, to be methylated by Bik3 to bikaverin. A haploid wild type S288C strain (WT-BIK) was additionally adaptively evolved.

**Figure 1.**
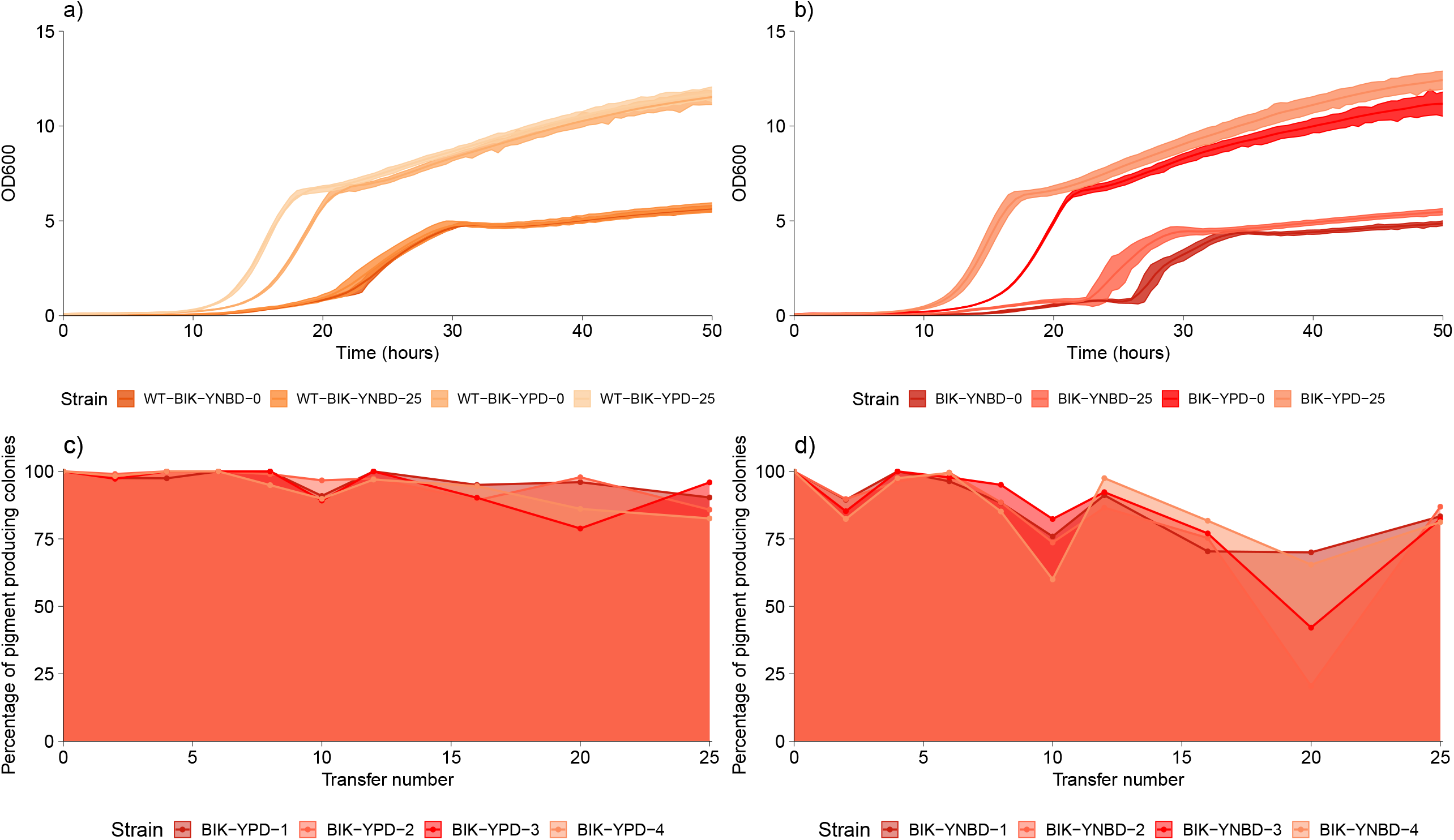
Experimental workflow. Four replicate lineages per strain, initiated from four *S. cerevisiae* CEN.PK strains (indigoidine producing haploid and diploid) and two *S. cerevisiae* S288C strains (wild type and bikaverin producing), were serially passaged in rich and synthetic defined media with galactose or glucose as a carbon source for 175-200 generations (up to 25 transfers) (a). Counting of the proportion of pigmented colonies was performed at regular intervals (b). Comparative growth characterization was performed for parental and evolved ALE end-point populations (c). Lastly, the parental strains and evolved end-point populations were whole genome sequenced to identify variants enriched during the ALE (d).

The four *S. cerevisiae* strains (IND-H, IND-D, WT-BIK, BIK) were each used to initialize four independent ALE lineages that were asexually adaptively evolved on galactose as the sole carbon source in rich medium (YPG) for ∼200 generations and in synthetic defined medium with ammonium as the nitrogen source (YNBG) for ∼175 generations. Galactose represented a favored, mostly respirative, carbon source for CEN.PK-based but unnatural for S288C-based strain that is a poor galactose utilizer. Galactose was also known to benefit the production of indigoidine by maintaining the cells’ metabolic state mostly respirative (Wehrs et al., 2018).

The CEN.PK-based lineages (originating from the parentals IND-H, and IND-D) grew efficiently on the galactose media from the first transfer whereas, as expected (Otero et al., 2010), the S288C-based lineages (originating from the parentals WT-BIK, and BIK) exhibited a long lag-phase of up to 46 h, on the YNBG medium in particular (Figures 2 and 3, Supplementary material, Table S1). The burden imposed by the heterologous pigment production was apparent in the growth profiles of the S288C wild type (WT-BIK) and bikaverin producing (BIK) parental strains. BIK parental strain had an almost twice as long lag phase and a significantly lower maximum specific growth rate (i.e., 0.19 h^-1^ on YPG and 0.15 h^-1^ on YNBG) than the S288C wild type (WT-BIK) strain (i.e., 0.36 h^-1^ on YPG and 0.25 h^-1^ on YNBG) (ANOVA and Tukey’s test, n = 3, P value < 0.01) (Figure 3, Supplementary material, Tables S1 and S2). Similarly, the maximum specific growth rates of the CEN.PK indigoidine producing (IND) haploid (i.e., 0.23 h^-1^ on YNBG and 0.38 h^-1^ on YPG) and diploid parental strains (i.e., 0.28 h^-1^ on YNBG and 0.40 h^-1^ on YPG) were lower than those of the CEN.PK wild type haploid (i.e., 0.33 h^-1^ on YNBG and 0.52 h^-1^ on YPG) and diploid (i.e., 0.21 h^-1^ on YNBG and 0.50 h^-1^ on YPG) strains had (ANOVA and Tukey’s test, n = 3, P value < 0.01) (Figure 2). The CEN.PK wild type and indigoidine producing parental strains had similar lag-phase lengths.

**Figure 2.**
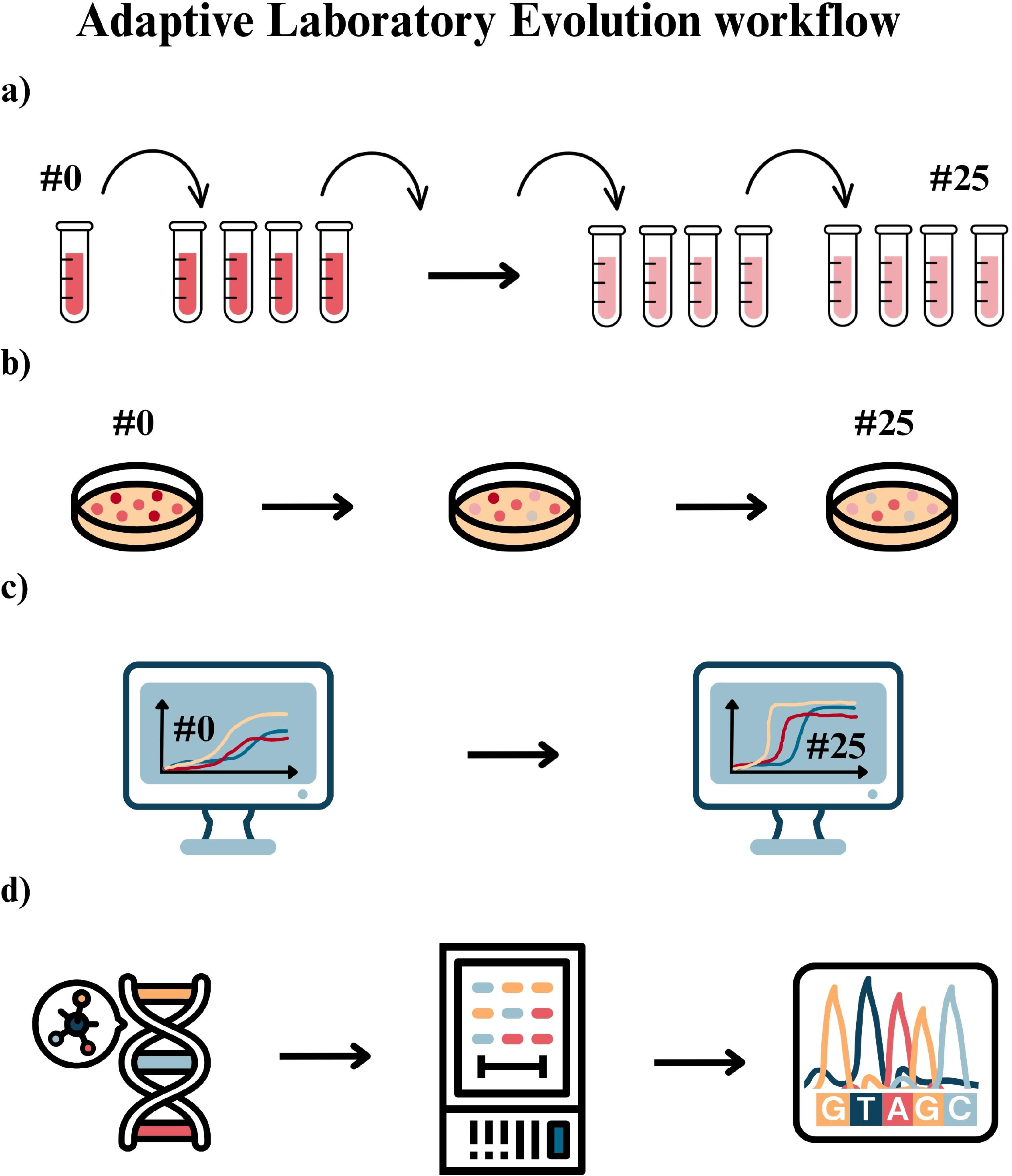
Growth profiles of haploid and diploid *S. cerevisiae* CEN.PK-based heterologous indigoidine pathway having parental strains IND-H (a) and IND-D (b) and evolved populations. The growth curves are shown as loess fits of ten biological replicate cultures for the parental strains (IND-H-YNBG-0, IND-D-YNBG-0, IND-H-YPG-0, IND-D-YPG-0). The growth curves for evolved ALE end-point populations are shown as loess fit averages of each of the four replicate lineages, with ten biological replicate cultures per lineage (IND-H-YNBG-25, IND-D-YNBG-25, IND-H-YPG-25, IND-D-YPG-25). The maximum growth rates of each biological replicate are shown for all c) IND-H-YPG and d) IND-H-YNBG parental and end-point populations. The growth profiles with significantly improved maximum specific growth rate compared to the parental are marked with stars (ANOVA and Tukey’s test; n=10; P value < 0.01).

**Figure 3.**
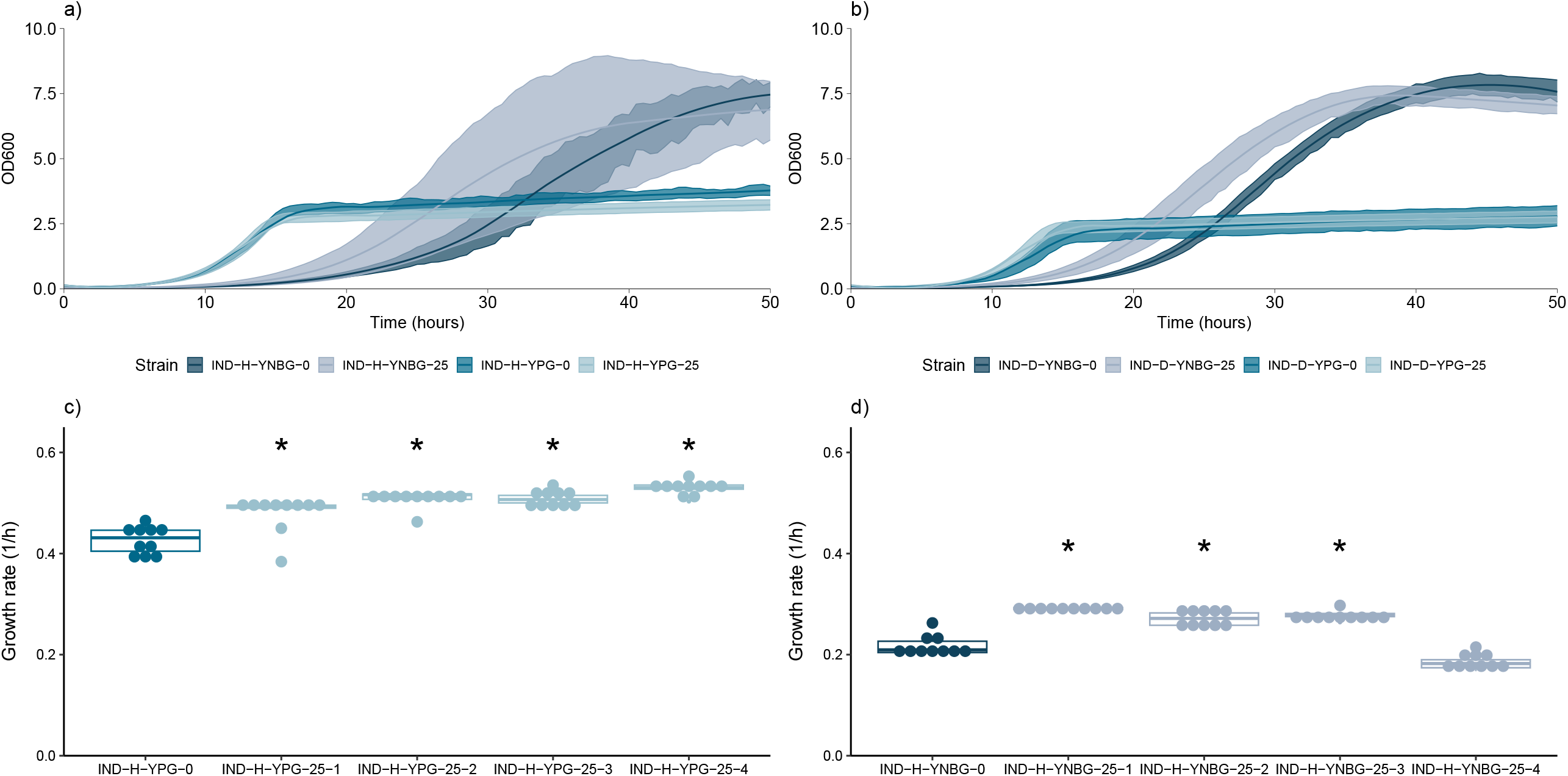
Growth profiles of haploid wild type *S. cerevisiae* S288C-based and heterologous bikaverin pathway having *S. cerevisiae* S288C-based parental strains (a) WT-BIK, b) BIK) and evolved populations. The growth curves are shown as loess fits of ten biological replicate cultures for the parental strain. The growth curves for the ALE end-point populations are shown for each of the four replicate lineages as loess fits of ten biological replicates per lineage. The maximum growth rates of each biological replicate are shown for all c) BIK-YPG and d) BIK-YNBG parental and end-point populations. The growth profiles with significantly improved maximum specific growth rate compared to the parental strain are marked with stars (ANOVA and Tukey’s test; n=10; P value < 0.01).

Comparative growth characterization of the parental and evolved populations (after 25 transfers and ∼175 to ∼200 generations depending on the medium) showed that most of the evolved S288C-based populations with bikaverin pathway (BIK-YPG-25-1 to 4, and BIK-YNBG-25-3 and 4) had notably improved maximum specific growth rates compared to their respective parental populations (i.e., an average 1.7 and 1.6 times faster in populations evolved on YPG and YNBG, respectively) (Figure 3, Supplementary material, Table S2). In addition, lag-phases were shortened nearly to half in S288C-based ALE lineages with bikaverin pathway compared to the respective parental populations independent of the evolution medium. In contrast, the wild type S288C -based evolved lineages had slightly faster (i.e., 1.2 times) maximum specific growth rates on YNBG but shorter (i.e., on average 12 h) lag-phase on YPG than the corresponding parental population (Figure 3). The lag-phases were not substantially shortened in evolved CEN.PK -based lineages with indigoidine pathway (Figure 2, Supplementary material, Table S3). In addition, all except one of the IND-based lineages grew faster, but not notably faster, than the respective parental populations (i.e., on average IND-H-YNBG-25-1 to 3: 1.3 times faster, IND-H-YPG-1 to 4: 1.3 times faster, IND-D-YNBG-1 to 4: 1.1 times faster, IND-D-YPG-1 to 4: 1.4 times faster) (Figure 2, Supplementary material, Table S4).

### Cell viability improved in indigoidine producing lineages

Since lineages having indigoidine pathway showed only little improvement in growth dynamics and the indigoidine pigment has been found to form sharp crystals that may burst the producer cells (Ghiffary et al., 2021), the cell viabilities of evolved populations from IND-H and IND-D ALE lineages and the corresponding parentals were assessed. The proportion of viable cells was found always higher on YPG than on YNBG medium. All except one of the lineages adaptively evolved on YNBG had developed improved cell viability during ALE (ANOVA and Tukey’s test, n = 3, P value < 0.01) (Figure 4, Supplementary material, Table S5). In addition, the haploid *S. cerevisiae* CEN.PK lineages with indigoidine pathway adaptively evolved on YPG medium improved their cell viability during ALE (ANOVA and Tukey’s test, n = 3, P value < 0.01) and one of the diploid ALE lineages (IND-D-YPG-25-3) could be detected to have a notably (ANOVA and Tukey’s test, n = 3, P value < 0.01) reduced cell death rate.

**Figure 4.**
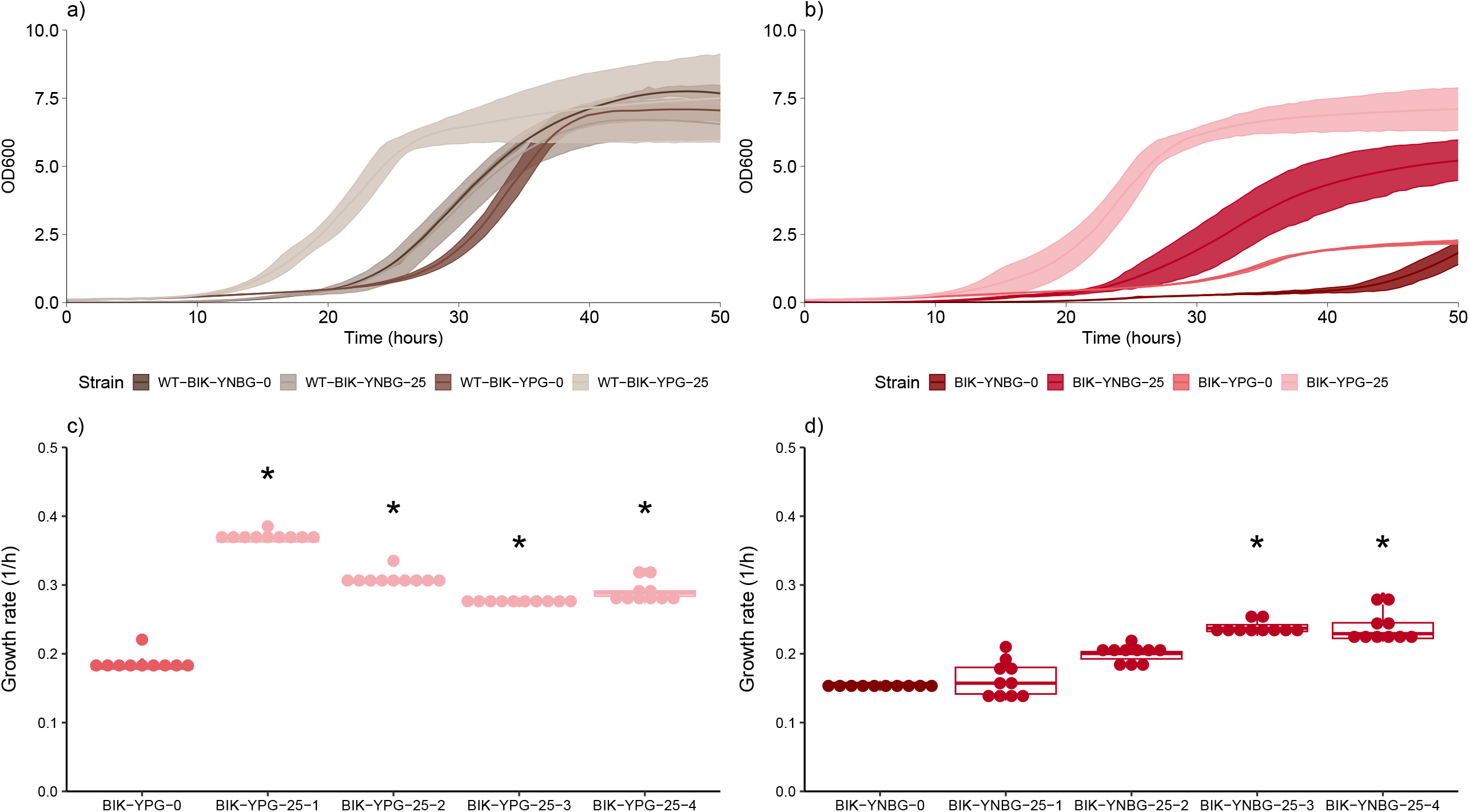
Percentage of cell death in parental and adaptively evolved heterologous indigoidine synthesis pathway having *S. cerevisiae* CEN.PK-based populations. IND-H and IND-D populations were haploid and diploid, respectively. The cell viability assessment was performed for cells grown in their respective adaptive evolution media, i.e., either in synthetic defined medium with galactose and ammonium as the carbon and nitrogen sources (YNBG) or on rich medium with galactose as the carbon source (YPG). The assayed populations included parental (0) and populations of four adaptive evolution lineages after 25 transfers (25-1, 25-2, 25-3, 25-4). The data represents the means and standard deviation from triplicate measurements. ^*^P < 0.01 indicates a significant difference compared to the parental strain using ANOVA and Tukey’s test (n = 3).

### Proportion of indigoidine pigmented colonies declined rapidly during ALE

To assess the stability of pigment production in adaptively evolving engineered strains, the proportion of pigmented colonies on plates were regularly determined (Figures 5-7). For all the ALE lineages initiated from pigment producing parental strains, colony counting was performed for each transfer 0 – 10 and then for every fifth transfer until the end of the ALE experiment. The individual colonies were counted and categorized into two separate groups: pigmented or non-pigmented. In both the haploid and diploid indigoidine pathway having CEN.PK-based lineages, pigmentation reduced rapidly on both YPG and YNBG media (Figure 6, Supplementary material, Figure S1). The proportion of indigoidine pigmented colonies reduced gradually in the YPG grown lineages, whereas on YNBG medium, the loss was quite sudden. Regardless of ploidy, the lineages remained predominantly strongly pigmented up to ∼5 transfers in YNBG and ∼10 transfers in YPG. In YNBG medium a major loss of pigmented colonies occurred by the transfer 10 whereas in YPG medium the proportion of pigmented colonies dropped in some of the lineages only by 20 transfers. By transfer 20, the slightly pigmented colonies showing shades of grey replaced the strongly pigmented deep blue colonies completely and non-pigmented colonies made up the majority (Figure 5). By the end of the ALE experiment (transfer 25) all except one (i.e., IND-D-YNBG-3) replicate lineage had lost all pigmented colonies.

**Figure 5.**
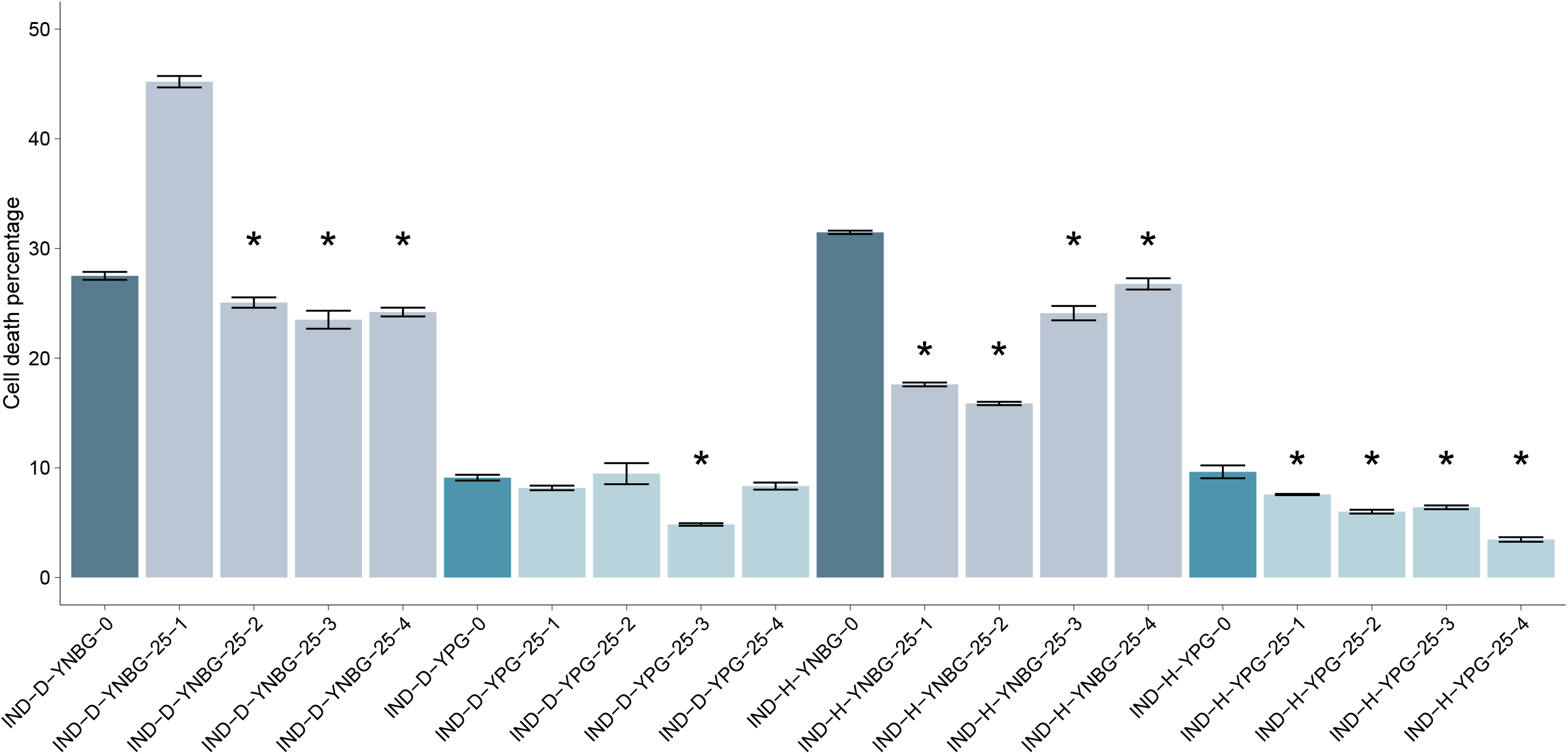
Colony pigmentation of indigoidine pathway having *S. cerevisiae* CEN.PK-based ALE lineages, adaptively evolved in medium with galactose as the sole carbon source. a) Representative population after the first batch culture in synthetic defined galactose medium (IND-D-YNBG-0), b) representative population after five transfers adaptively evolved in rich galactose medium (IND-D-YPG-5), c) representative population after ten transfers adaptively evolved in rich galactose medium (IND-H-YPG-10), d) representative population after fifteen transfers adaptively evolved in rich galactose medium (IND-D-YPG-15), e) representative population after 20 transfers adaptively evolved in synthetic defined galactose medium (IND-H-YNBG-20), and f) representative ALE end-point population adaptively evolved in synthetic defined galactose medium (IND-H-YNBG-25). The images show the decrease of highly pigmented colonies and the emergence of slightly pigmented and non-pigmented colonies during the progression of ALE.

**Figure 6.**
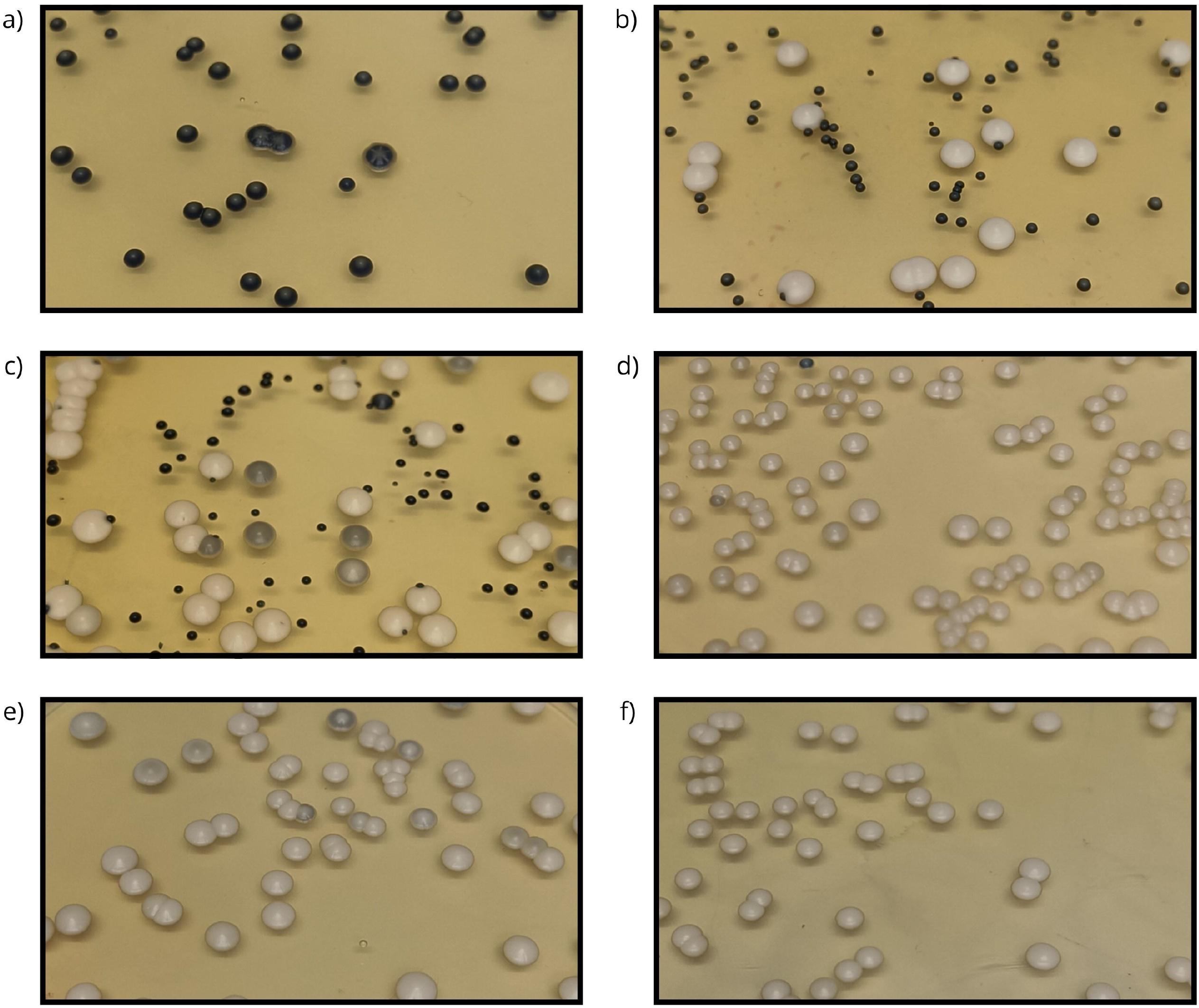
Percentage of pigmented colonies as a function of transfers in the ALE experiment in lineages having indigoidine pathway (a-b) and in lineages having a bikaverin pathway (c-d). Colony counting of all and pigmented colonies was performed for each ALE lineage on single plates except for the ALE end-point population in four replicates for each lineage.

**Figure 7.**
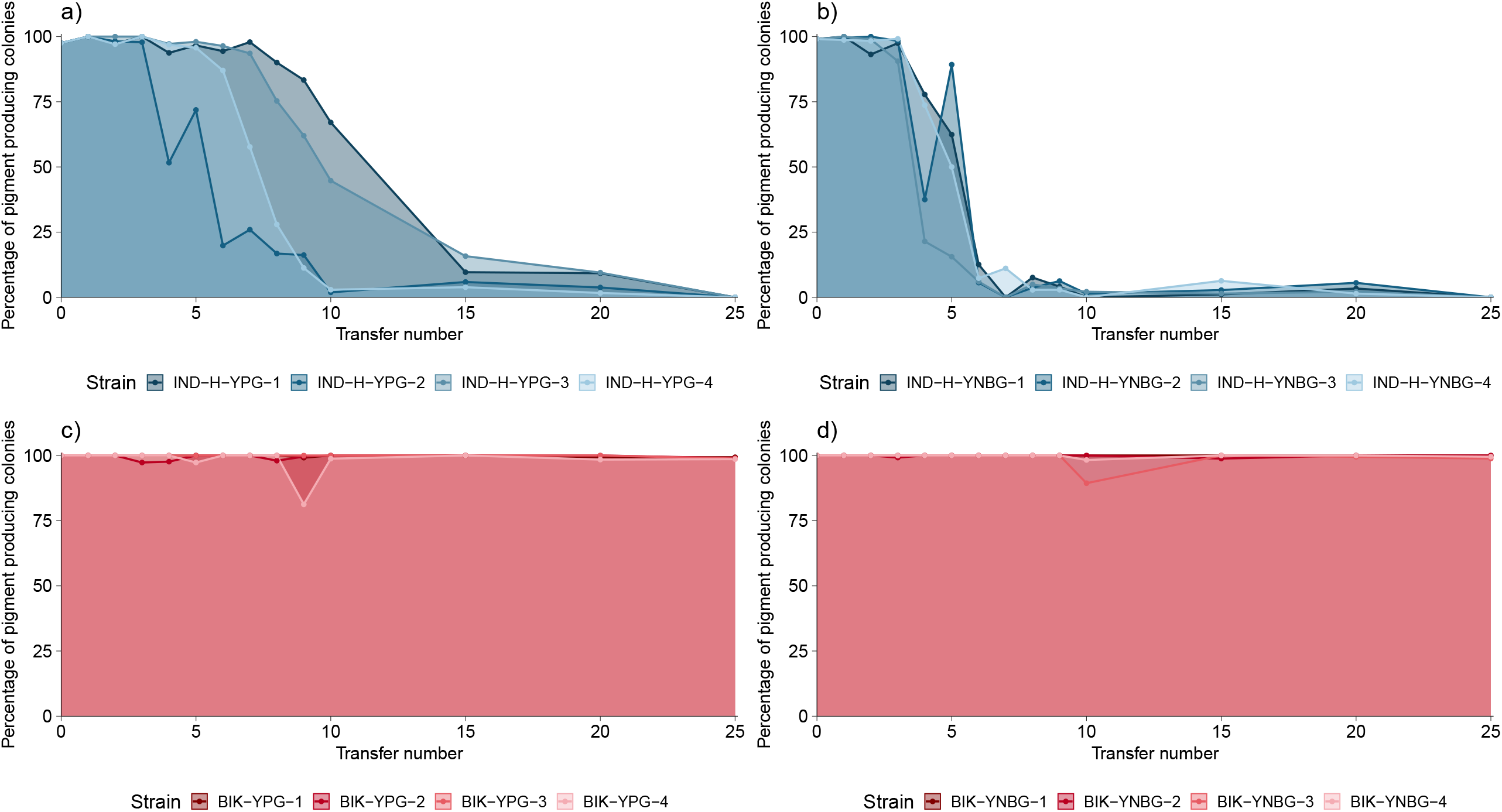
The robust colony pigmentation and intensity differences of bikaverin pathway having haploid *S. cerevisiae* S288C ALE lineages. Representative populations after the first batch culture in rich galactose medium (BIK-YPG-0) (a), after ten transfers in synthetic defined galactose medium (BIK-YNBG-10) (b), after 20 transfers in synthetic defined galactose medium (BIK-YNBG-20) (c), at the end of ALE in rich galactose medium (BIK-YPG-25) (d), at the end of ALE in rich glucose medium (BIK-YPD-25) (e), and at the end of ALE in synthetic defined glucose medium (BIK-YNBD-25) (f).

### Bikaverin pigmentation was notably stable in ALE on galactose

In contrast to the CEN.PK-based lineages having indigoidine pathway (IND-H and IND-D), the S288C-based lineages having bikaverin pathway (BIK) remained almost fully pigmented throughout the ALE experiment (Figure 6 and 7). In later populations (i.e., from transfer 10 onwards), individual colonies had the most intense colour in the centre of the colony that faded towards the colony edges (Figure 7, Supplementary material, Figure S2). In the S288C-based lineages the majority of the colonies remained strongly pigmented until the end of the ALE experiment on the YNBG medium (Figure 6 and 7), and those on YPG medium also maintained over 90% proportion of highly pigmented dark red colonies, though less intensely red than on YNBG medium.

### During ALE on glucose lag-phases were reduced

To assess whether bikaverin production was equally robust on a favored chemical environment, the wild type and bikaverin producing S288C strains (WT-BIK, BIK) were used to initiate ALE lineages on glucose as the carbon source in rich (YPD) and synthetic defined (YNBD) media. During the ALE, the maximum specific growth rates did not improve (Supplementary material, Table S6) but the lag-phases slightly shortened in most of the lineages compared to the respective parental populations (i.e., at most by 4 h on average in BIK lineages on YPD) (Figure 8, Supplementary material, Table S7). The lineages with bikaverin pathway (BIK) adaptively evolved on glucose remained predominantly pigmented throughout the ALE like the BIK lineages adaptively evolved on galactose media (Figure 8). However, in particular the BIK lineages evolved on YNBD enriched more non-producers (Figure 8) than the BIK lineages evolved on galactose as the carbon source (Figure 6). Qualitatively the lineages evolved on glucose showed increasingly many faintly pigmented colonies by the end of ALE, compared to the robust intense pigmentation observed even in ALE end-point populations on galactose (Figure 7).

**Figure 8.**
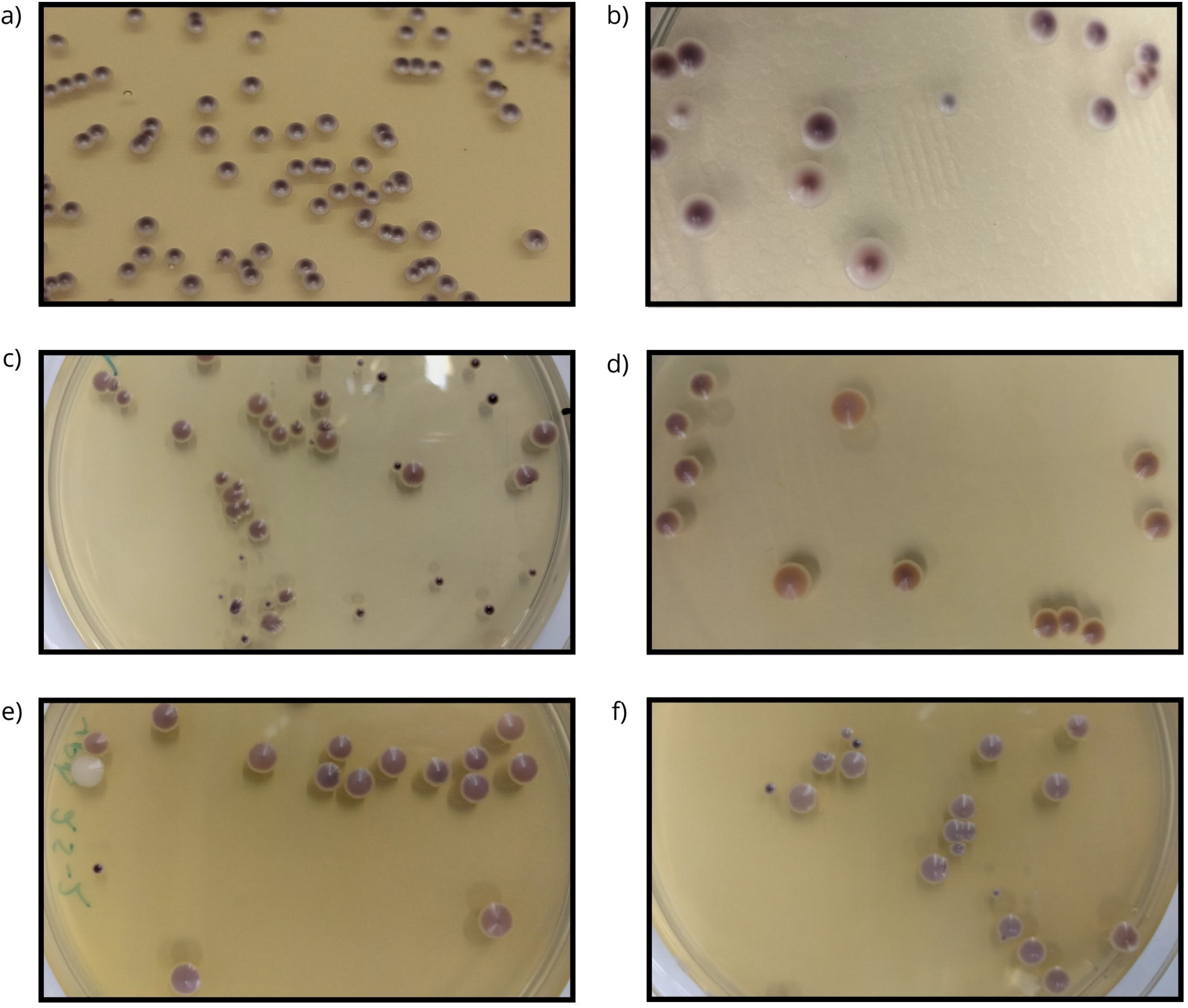
Growth profiles of the haploid wild type (WT-BIK) (a) and bikaverin pathway having (BIK) (b) *S. cerevisiae* S288C parental and evolved populations. The growth curves are shown as loess fits of ten biological replicates for the parental strain. The growth curves for the ALE end-point populations are shown for each of the four replica lineages as loess fits of ten biological replicates per lineage. Percentages of bikaverin pigmented colonies as a function of transfers in the ALE experiments in rich glucose medium (YPD) (c) and in synthetic defined glucose medium (YNBD) (d). Colony counting of all and pigmented colonies was performed for each ALE lineage on single plates except for the ALE end-point population in three replicates for each lineage.

### Adaptive solutions included mutations in heterologous genes or in genes involved in sugar utilization

To identify mutations that enriched in the lineages during ALE, most initial populations (transfer 0) and ALE end-point populations (transfer 25) were whole-genome sequenced. In CEN.PK-based lineages with indigoidine pathway mutations were detected in indigoidine synthetase BpsA. A frequent hot spot single nucleotide deletion in BpsA encoding gene was detected across diploid lineages evolved on YNBG (pos.1968, AT->T, VAF 58% to 78%) (Supplementary material, Table S8). The mutation leads into a frameshift and, thus, it was likely a loss of function mutation. Less frequent hot spot single nucleotide variants (SNVs) in the BpsA encoding gene were detected across different combinations of haploid lineages evolved on YNBG (pos. 1346 G->T, variant allele frequency (VAF) 18% to 33%, and either pos.255 G->A, VAF 12% to 22%, or pos.1831 C->A, VAF 10% to 13%). In contrast, no missense, frameshift, or truncating variants were detected in the coding sequences of native genes above VAF 20% in CEN.PK-based ALE lineages with indigoidine pathway.

In S288C-based lineages with bikaverin pathway missense variants were recurrently detected in the native *GAL2* gene independent of the medium (BIK-YNBG-25-1, p.Thr368Ala, VAF 63%; BIK-YNBG-25-2, p.Cys77Arg, VAF 92%; BIK-YNBG-25-4, p.Leu74His, VAF 99%; BIK-YPG-25-1, p.Ser369Tyr, VAF 99%; BIK-YPG-25-2, p.His392Arg, VAF 65%; and BIK-YPG-25-3, p.His392Arg, VAF 99%) (Supplementary material, Table S9). *GAL2* encodes a galactose permease, which is required for the utilization of galactose but found non-functional in S288C wild type (Mortimer and Johnston, 1986; Otero et al., 2010). BIK-YNBG-25-1 and 4 had also missense mutations in a transcription factor encoding *RGT1* (p.Asp781Tyr, VAF 98%, and p.Asp781Tyr, VAF 98%, respectively). In an absence of glucose Rgt1 acts as a repressor of genes involved in sugar utilization which is likely a function affecting cells’ fitness on galactose (Kim et al., 2003; Özcan et al., 1996). In BIK-YNBG-25-1 population a variant was detected also in hexose transporter encoding *HXT10* (p.Ala146Val, VAF 81%) and in BIK-YNBG-25-4 a variant in *BUR2* (p.Leu300Pro, VAF 75%). BIK-YPG-25-1 and BIK-YPG-25-2 had enriched truncating variants in *WHI2* (p.Arg365*, VAF 99%, and p.Glu31*, VAF 66%, respectively) encoding a suppressor of TORC1 in low amino acid abundance (Chen et al., 2018). BIK-YPG-25-3 had instead two high frequency variants in polyubiquitin-binding protein encoding *DSK2* (p.Ala123Ser, VAF 100%) and in *NOB1* (p.Glu248Lys, VAF 81%) involved in proteasomal and 40S ribosomal subunit biogenesis. No bikaverin pathway variants were detected in alternative allele frequency higher than 7%. Thus, bikaverin pathway having S288C-based lineages from both YNBG and YPG had enriched SNVs mainly in genes involved in sugar utilization, but variants detected in the heterologous genes were rare. In contrast, the CEN.PK-based ALE lineages with indigoidine pathway had enriched variants mainly in heterologous indigoidine synthetase.

On glucose evolved S288C-based lineages with bikaverin pathway had enriched mutations in few native genes. Recurrently mutated were *MHP1* involved in microtubule organization in lineages evolved on YNBD (BIK-YNBD-25-1, p.Glu571*, VAF 73%, BIK-YNBD-25-3, p.Trp22*, VAF 51%, BIK-YNBD-25-4, p.Gly155fs, VAF 97%) and *PNO1* involved in pre-18S rRNA processing in lineages evolved on YPD (BIK-YPD-25-1, p.Arg105Lys, VAF 74% and BIKYPD-25-4, p.Gly203Arg, VAF 58%) (Supplementary material, Table S9). However, bikaverin pathway variants were detected in S288C-based lineages evolved on YPD up to variant allele frequency 14.2% and in populations evolved on YNBD up to variant allele frequency of 12%. Thus, the frequences of bikaverin pathway variants were on the similar level to the percentages of colonies having lost the pigmentation (Figure 8).

## Discussion

Cells with engineered pathways are likely to have a lower fitness than their evolutionarily streamlined wild type counterparts. Here, engineered pigment producing strains manifested fitness compromises as reduced maximum specific growth rates and cell viability and extended lag-phases compared to the respective background strains. These are common consequences of protein over-expression (Eguchi et al., 2018; Fujita, 2025; Krogh et al., 2008) Due to such fitness-compromises the engineered traits are exposed to negative Darwinian selection in the presence of non-producer clones arising from DNA-replication associated random mutations. Previously, Darwinian selection has been observed to decline the production of N-acetylglucosamine in *S. cerevisiae* (i.e., within 150 – 200 generations) (Lee et al., 2021) and, more rapidly, the production of vanillin-β-glucoside (i.e., decline up to 91% of production from 14 to 35 generations) (D’Ambrosio et al., 2020). Since evolutionary adaptation of microbial cells is commonly not limited by the mutation rate (Arjan G et al., 1999; Hall et al., 2010), the rate of production loss is likely to reflect the relative fitness benefit. Here, indigoidine production declined rapidly but bikaverin production was highly robust over the course of generations, though qualitatively reduced in intensity. However, the galactose media posed an environmental challenge to the bikaverin producing gal2-S288C strain (Mortimer and Johnston, 1986), but not to the indigoidine producing CEN.PK strains. The fitness benefit from adapting to the environmental challenge appeared to be notable, since the S288C-based ALE lineages with bikaverin pathway adapted up to 40% higher maximum specific growth rate and shorter lag-phase without a notable pigmentation decline. Previously, 24% faster growing *S. cerevisiae* CEN.PK113-7D strains were obtained after ALE on galactose (Hong et al., 2011). On glucose the expected achievable fitness benefit from an adaptation to the environment was low independent of the background strain. Previous long-term ALE of *S. cerevisiae* S288C descendant W303 (not gal2-) on galactose and glucose found a fitness benefit from adapting to the serial culturing on glucose of 3.7⍰±⍰0.06%, which was less than half of that on galactose (i.e., 8.2⍰±⍰0.1%) (Martinez et al., 2023). Correspondingly, when the adaptation to the environment offered a low fitness benefit, bikaverin pigmentation decline was observed during ALE on glucose. Thus, relative fitness-benefit achievable from an adaptation to the growth environment is determining in the robustness of engineered traits.

Since the occurrence of beneficial mutations (if they are not rare) appears not to limit adaptation (Arjan G et al., 1999; Hall et al., 2010), the mutational target sizes of traits are not likely to affect their relative robustness. However, could the mutational target size of a production trait be so small that mutations disrupting the trait would be rare? Here, the indigoidine and bikaverin production traits differed in the mutational target size though the exact sizes were not determined. Indigoidine production trait is dependent on two enzymes, both of which are required for the pigment synthesis. Similarly, Bik1 and npgA activities are necessary for pre-bikaverin synthesis. However, pre-bikaverin has weaker color than the later pathway products, and therefore disruptions also in Bik2-Bik3 lead to pigmentation loss.

Several means such as growth-product coupling and product addiction, have been proposed for increasing the robustness of engineered traits (Lv et al., 2020; Rugbjerg et al., 2018b). These involve expression of synthetic regulatory components but commonly also metabolic gene deletions. Compensatory evolution in *S. cerevisiae* in response to gene deletions has been found rapid and cover a wide variety of trait defects (Szamecz et al., 2014). Accordingly, propagating engineered cells having e.g., growth-product coupling, optimized promoters, and inactivated competing pathways have been reported to revert to wild type phenotypes restoring cellular fitness (D’Ambrosio et al., 2020; Jacobsen and Frigaard, 2014; Pereira et al., 2021; Yunus and Jones, 2018). Thus, the challenge of production loss due to fitness compromises of engineered traits is yet unresolved. Findings from this work suggest that in order to ensure the robustness of production strains, the fitness compromises and mutational target sizes of the engineered traits including any synthetic regulatory features should be assessed in the strain development in the context of the application chemical environment and controlling the relative fitness benefits may be the key.

## Material and Methods

### Media and strain construction

For strain construction, *E. coli* containing relevant plasmids were grown in lysogeny broth (LB) with 100 µg/ml ampicillin (MERCK). For transformations, cells were cultivated on YPD containing 20 g/L of bacteriological peptone (Neogen), 10 g/L of yeast extract (Neogen), and 20 g/l D-glucose (VWR Chemicals). For selection and maintenance of plasmids after and during transformation, selective YPD media with 200 µg/ml nourseothricin dihydrogen sulfate (NAT) (Jena Bioscience, AB-101) and/or 200 µg/ml Geneticin® G-418 sulfate powder (G418) (MERCK) was used. For solid media 20 g/L of bacteriological agar (VWR Chemicals) was used with relevant plate compositions. Solid YPD was also used for the colony counting, to reduce the potential effect of phenotypical adaptation on the colony pigmentation.

All *S. cerevisiae* transformations were performed using the standard lithium acetate protocol (Gietz, 2014) and the CRISPR/Cas9 protocol of the EasyClone kit (Jessop-Fabre et al., 2016), with expression plasmids linearized by NotI enzyme (FD0596, Thermo Scientific). Correct integration was confirmed with QuickLoad Taq PCR (NEB).

ALE, growth characterizations and genomic DNA extraction were performed using media with galactose or glucose as the carbon source: rich media (YPG and YPD) and synthetic defined media (YNBG and YNBD). The media contained either 30 g/L D-(+)-galactose (Sigma-Aldrich) or 20 g/l D-glucose (VWR Chemicals) as carbon source. The rich medium contained 10 g/L of yeast extract (Neogen), and 20 g/L of bacteriological peptone (Neogen) (YP). The synthetic defined medium contained 6.7 g/L yeast nitrogen base without amino acids (YNB, Sigma-Aldrich). The yeast nitrogen base without amino acids contained 5.0 g/L of ammonium sulfate as its nitrogen source, 2.0 mg/L of inositol, 400 μg/L of calcium pantothenate, 400 μg/L of pyridoxine HCl, 400 μg/L of nicotinic acid, 400 μg/L of thiamine HCL, 200 μg/L of p-aminobenzoic acid, 200 μg/L of riboflavin, 2.0 μg/L of folic acid, and 2.0 μg/L of biotin as vitamins. The trace elements included 500 μg/L of boric acid, 400 μg/L of zinc sulfate, 400 μg/L of manganese sulfate, 200 μg/L of ferric chloride, 200 μg/L of sodium molybdate, 100 μg/L of potassium iodide and 40 μg/L of copper sulfate. The salts included 1.0 g/L of potassium phosphate monobasic, 0.5 g/L of magnesium sulfate, 0.1 g/L of calcium chloride and 0.1 g/L of sodium chloride.

### Strains

All the strains constructed and referenced in this study are listed in Supplementary material, Table S10. The indigoidine production pathway was engineered in a haploid *S. cerevisiae* CEN.PK113-7D strain and a diploid cross between CEN.PK113-7D and CEN.PK113-1A. The haploid wild type strains CEN.PK113-1A and CEN.PK113-7D were provided by Euroscarf. The haploid bikaverin producing *S. cerevisiae* S288C strain and wild type *S. cerevisiae* S288C strain were kindly provided by Prof. Uffe Hasbro Mortensen (Technical University of Denmark, Denmark). Both the bikaverin production pathway and the indigoidine production pathway were assembled at EasyClone locus XII-5 (Mikkelsen et al., 2012). The haploid indigoidine and bikaverin producing strains had the nourseothricin (NAT) antibiotic resistance marker integrated. The indigoidine diploid strains had *PEP4* and *PRB1* deleted with a double deletion. All modified strains had Cas9 expressing plasmids containing the antibiotic resistance marker for G418 (Geneticin) removed before experimental culturing was conducted.

### Adaptive laboratory evolution

Each of the six strains (WT-H, WT-D, IND-H, IND-D, WT-BIK, BIK, Supplementary material, Table S10) was inoculated in YNBG and YPG media and WT-BIK and BIK strains also in YNBD and YPD to create the parental populations (transfer 0). Cell mass was taken from solid agar plates from multiple different colonies to increase the initial genetic variation, with only pigment producing colonies being chosen for the indigoidine and bikaverin producing strains.

From the parental cultures (transfer 0), four parallel lineages were inoculated per medium into a starting OD of 0.1 (transfer 1). The CEN.PK strains had a total of 8 total lineages per strain, 4 YPG medium and 4 in YNBG medium. The S288C strains had a total of 16 lineages per strain, 4 in YPG medium, 4 in YNBG medium, 4 in YPD medium and 4 in YNBD medium. Each of the cultures was grown in 4 ml of their respective media in 12 ml round bottom culture tubes. Cells were grown for 2-3 days at 30°C with 220 rpm shaking in the Innova 44 Incubator Shaker (New Brunswick). From the second transfer (#2) onwards, 1% (40 μL) of the cultures in rich medium (i.e., YPG or YPD) were transferred to fresh medium, while 2% (80 μL) of the cultures in synthetic defined medium (i.e., YNBG or YNBD) were transferred. For transfers 8 and 9 of the haploid indigoidine producing strain, 100 mg/L ampicillin was added to all cultures to remove bacterial contamination. Bacterial contamination was not observed afterwards or in any other cultures. All ALE lineages were transferred up to end-point populations (transfer 25).

Glycerol stocks of each lineage were saved at -80°C for all transfers 0-10. From then onwards, glycerol stocks of the lineages at transfers 15, 20, and 25 were saved. For glycerol stocking, 300 µl of 50% glycerol was diluted with 700 µl of each cell culture for a final percentage of 15% glycerol per tube.

### Colony counting

To track the production stability, pigment producing populations were diluted and plated on solid YPD for colony counting. The total number of colonies, number of pigmented colonies, and level of pigmentations was assessed visually. Colony counting was performed for all transfers 0-10 of the indigoindine and bikaverin pathway having lineages. From then onwards, colony counting was performed at transfers 15, 20, and 25. At transfers 0-10, 15, and 20, cell cultures were diluted and plated in a single replicate. At transfer 25, four replicate plates per cell culture were prepared from the same dilution. Dilution series of each culture tube was prepared prior to plating. A dilution of 1/10000, 1/100000, or 1/1000000 was plated, depending on the growth medium to achieve a suitable number of colonies on the plates (40-700). The 1/10000 dilution factor was used in the beginning of the ALE experiment, being completely replaced by the 1/100000 dilution factor after a couple of transfers for the cultures in synthetic defined medium and 1/1000000 for the cultures in rich medium. 100 µl of diluted cell culture was evenly spread using an L-shaped spreader and the plates were incubated at 30°C for 2–3 days, after which they were left at 4°C. Colonies were counted within a week of placing the plates in the cold for the bikaverin lineages and within 2-3 weeks for the indigoidine lineages. Bikaverin pigmentation of colonies developed without a delay whereas indigoidine pigmentation developed more slowly. For the numerical representation of colony pigmentation, no distinction was made regarding the pigment intensity.

### Growth characterization

For growth characterization, initial populations (transfer 0) and end-point populations (transfer 25) of all ALE lineages were cultivated in their respective media. 4 ml of the respective media, rich or synthetic defined, in 12 ml tubes were inoculated from cryo stocks and incubated at 30°C with 220 rpm shaking in the Innova 44 Incubator Shaker (New Brunswick) for 4-5 days which was a compromise between parental and evolved populations. The parental populations WT-D-YPG, IND-D-YPG, BIK-YNBG did likely not reach stationary phase in these precultures. For OD measurements, cells were cultured in 100-well flat-bottom honeycomb plates with 150 µl respective medium inoculated into a start OD of 0.02 in ten biological replicates. The cultures were incubated and the turbidities monitored using Bioscreen FP-1100-C (Labsystems) at 30°C, with continuous shaking, high amplitude and fast speed. The 600 nm brown filter was used for turbidity measurements every 30 minutes. The Bioscreen data was transformed using ODtransformed = OD + 0.8324 * OD^3^ (Warringer et al., 2003) and converted to corresponding spectrophotometer turbidity values by taking into account the light path length of 0.39 cm when the Bioscreen well radius was 0.35 cm. Lag phase lengths were estimated according to (Warringer and Blomberg, 2003).

### Growth rate calculations and statistical analysis

The growth rates were calculated using R v. 4.3.2 with package growthcurver v. 0.3.1 (Sprouffske and Wagner, 2016). The statistical significance of the growth rate changes was assessed with ANOVA and Tukey’s test.

### Genomic DNA extraction

Cells were cultivated in 4 ml of their respective ALE mediums for genomic DNA extraction. Cultures were inoculated with a loop from a glycerol stocks and incubated at 30°C with 220 rpm shaking in 12 ml culture tubes. The ALE end-point populations (transfer 25) were incubated for 2-3 days and the parental strains (transfer 0) were incubated for 2-9 days to accumulate sufficient biomass for the genomic DNA extraction. Genomic DNA was extracted from the pellets using the MasterPure™ Yeast DNA Purification Kit (MPY80200 Lucigen, BioSearch Tech) according to manufacturer’s protocol with certain modifications. Instead of adding RNase during the initial 65°C incubation, 5 µl RNase was added before the isopropanol step, with incubation for 1 h at 37°C, to get rid of RNA contamination. An additional ethanol evaporating step, for 30 s at 60°C, was added after ethanol washing, to remove impurities as efficiently as possible. An extra RNase step was added after initial purification if needed. The purified samples had 1.5 µl RNase added with a 30 min incubation at 37°C. The sample was then re-purified according to manufacturer’s protocol from the isopropanol step onwards, with the additional ethanol evaporation step included.

### Cell viability assay

For cell viability assay, all strains were pre-cultured from CRYO stocks in 50 ml respective medium in 250 ml flasks for 2 nights, 30°C, 220 rpm. The pre-cultures were used to inoculate the experimental cultures of 50 ml fresh medium in 250 ml into a starting OD600 of 0.1. The cultures were incubated for 3 days, at 30°C, with 220 rpm shaking. Cell viability assay was conducted immediately after the incubation, with final OD measured for all flasks (Supplementary material, Table S11). Yeast cells were stained using LIVE/DEAD FungaLight Yeast Viability Kit (Life Technologies) according to manufacturer’s instructions to distinguish healthy and cell wall compromised cells. In short, yeast cells from 1 ml of a shake flask culture were harvested and washed with PBS pH 7.4 once before being sonicated to remove cell clumps. Each sample was then diluted with the same buffer to the final OD of 0.2. The diluted samples were stained with PI and SYTO 9 fluorescent dyes and analyzed using BD FACSAria III (BD Biosciences). Measurement of each sample was stopped when 2000 events of dead cell were counted. Triplicate measurements were performed for each culture.

### Whole genome sequencing

Library preparations and NGS sequencing was performed by Novogene on Illumina NovaSeq X Plus Series platform (Microbial Whole Genome Sequencing (WOBI) producing PE150 reads). The raw reads of genomic DNA samples (Supplementary material, Table S12) were deposited in ENA database (https://www.ebi.ac.uk/ena/) in PRJEB100615.

### Whole genome sequence data analysis

The quality of the obtained reads was checked using Fastqc v. 0.12.0 (Andrews, 2010). Adapter removal and low quality read filtering was performed using cutadapt v. 4.9 (Martin, 2011). The trimmed reads were aligned to *S. cerevisiae* CEN.PK113-7D reference genome (GCA_002571405.2_ASM257140v2) (Salazar et al., 2017) or *S. cerevisiae* S288C reference genome (S288C_reference_sequence_R64-5-1_20240529) with the Burrows-Wheeler Aligner v. 0.7.17 mem (Li and Durbin, 2009) using default parameters. The alignments were processed (added read groups, sorted, reordered, and indexed) and duplicate reads were marked using Picard Tools v. 3.0.0 (Poplin et al., 2017; Van der Auwera et al., 2013). SNV and indel variant calling was performed against the parental populations used to inoculate the replicate ALE lineages with GATK4 v. 4.6.0.0 Mutect2 (Poplin et al., 2017; Van der Auwera et al., 2013) using default parameters. The variant calls were filtered using GATK4 v. 4.6.0.0 (Poplin et al., 2017; Van der Auwera et al., 2013) FilterMutectCalls using default thresholds. SnpEff v.5.1 (Cingolani et al., 2012) was used to annotate the filtered variants. Variants with high median mapping quality of ∼60 were assessed. The same pipeline was run separately for the heterologous genes of indigoidine pathway (Supplementary material, Table S13) and bikaverin pathway (Zhao et al., 2024).

### Data analysis and visualization

Data analysis and visualization was performed using R v. 4.3.2 and packages ggplot2 v. 3.5.1, wesanderson v. 0.3.7, RColorBrewer v. 1.1-3, cowplot v. 1.1.3, tidyverse v. 2.0.0.

## Data Availability

Raw whole-genome sequencing was stored in ENA project: PRJEB100615.

## Acknowledgements

Diogo Antunes and Hoa Chu are thanked for experimental assistance.

Paula Jouhten acknowledges funding from Research Council of Finland (decision number 352417) and Novo Nordisk Foundation (NNF22OC0080180). The authors acknowledge the computational resources provided by the Aalto Science-IT project.

## Author contributions

Natalia, Kakko von Koch: Formal analysis, Investigation, Visualization, Writing – original draft, Writing -review & editing; Olli, Lohilahti: Investigation, Visualization, Writing - review & editing; Andreas, Møllerhøj Vestergaard: Investigation, Writing - review & editing; An, Nguyen: Investigation, Writing - review & editing; Tomas, Strucko; Investigation, Writing - review & editing; Paula, Jouhten: Conceptualization, Funding acquisition, Supervision, Formal analysis, Investigation, Writing – original draft, Writing - review & editing.

## Supplementary material

**Table S1.** Lag phase lengths of the end-point populations of four replicate ALE lineages and respective parental populations of *S. cerevisiae* S288C wild type (WT) and bikaverin pathway having (BIK) lineages adaptively evolved on YPG and YNBG media and grown on the respective media.

| Lineage | Parental population lag length (h) | Evolved population lag length (h) | $\Delta h$ |
| --- | --- | --- | --- |
| WT-BIK-YPG | 28.0 | 15.5 | -12.5 |
|  |  | 17.6 | -10.4 |
|  |  | 16.8 | -11.1 |
|  |  | 15.5 | -12.4 |
| WT-BIK-YNBG | 23.6 | 22.8 | -0.8 |
|  |  | 26.7 | +3.1 |
|  |  | 24.5 | +0.9 |
|  |  | 22.5 | -1.1 |
| BIK-YPG | 34.2 | 20.0 | -14.2 |
|  |  | 18.9 | -15.3 |
|  |  | 14.2 | -20.0 |
|  |  | 20.0 | -14.2 |
| BIK-YNBG* | 46.2 | 28.7 | -17.5 |
|  |  | 24.0 | -22.2 |
|  |  | 27.5 | -18.7 |
|  |  | 25.7 | -20.5 |

\*Slow growing parental population preculture may not have reached stationary phase before inoculation. Thus, the lag phase should be considered minimum achievable.

**Table S2.** Maximum specific growth rate (rho) estimates of the end-point populations of four replicate ALE lineages and the respective parental populations of *S. cerevisiae* S288C wild type (WT) and bikaverin pathway having (BIK) lineages adaptively evolved on YPG and YNBG media and grown in multi-well plate format on the respective media. Tukey’s test P-values are shown for the differences between ALE end-point populations’ and parental population’s maximum rho (one-way ANOVA and Tukey’s test (n_control_ = 10, n_case_ = 10), P-value < 0.01).

| Lineage | Parental population<br>$\rho$ ($\text{h}^{-1}$) | Evolved<br>population<br>$\rho$ ($\text{h}^{-1}$) | Tukey's test<br>P-value |
| --- | --- | --- | --- |
| WT-BIK-YPG | $0.36 \pm 0.01$ | $0.36 \pm 0.01$ | 0.91 |
| | | $0.35 \pm 0.01$ | 0.42 |
| | | $0.30 \pm 0.04$ | $3.3\text{E-}09$ |
| | | $0.17 \pm 0.01$ | $2.4\text{E-}13$ |
| WT-BIK-YNBG | $0.25 \pm 0.01$ | $0.30 \pm 0.01$ | $2.1\text{E-}12$ |
| | | $0.32 \pm 0.01$ | $2.4\text{E-}13$ |
| | | $0.29 \pm 0.01$ | $1.3\text{E-}09$ |
| | | $0.30 \pm 0.01$ | $2.7\text{E-}12$ |
| BIK-YPG | $0.19 \pm 0.01$ | $0.39 \pm 0.01$ | $2.4\text{E-}13$ |
| | | $0.33 \pm 0.01$ | $2.4\text{E-}13$ |
| | | $0.29 \pm 0.01$ | $2.4\text{E-}13$ |
| | | $0.28 \pm 0.02$ | $2.4\text{E-}13$ |
| BIK-YNBG | $0.15 \pm 0.01$ | $0.15 \pm 0.03$ | 1.0 |
| | | $0.17 \pm 0.01$ | 0.023 |
| | | $0.23 \pm 0.01$ | $7.2\text{E-}13$ |
| | | $0.25 \pm 0.02$ | $2.5\text{E-}13$ |

**Table S3.** Lag phase lengths of the end-point populations of four replicate ALE lineages and respective parental populations of *S. cerevisiae* CEN.PK -based lineages with indigoidine pathway (IND) haploid (H) and diploid (D) lineages adaptively evolved on YPG and YNBG media and grown in multi-well plate format in the respective media.

| Lineage | Parental population lag length (h) | Evolved population lag length (h) | $\Delta h$ |
| --- | --- | --- | --- |
| IND-H-YPG | 11.8 | 11.1 | ND |
|  |  | 11.6 | ND |
|  |  | 12.0 | ND |
|  |  | 12.2 | ND |
| IND-H-YNBG | 24.1 | 18.5 | -5.6 |
|  |  | 18.3 | -5.9 |
|  |  | 16.9 | -7.2 |
|  |  | 32.4 | +8.3 |
| IND-D-YPG | 12.8 | 11.4 | ND |
|  |  | 11.7 | ND |
|  |  | 12.1 | ND |
|  |  | 11.2 | ND |
| IND-D-YNBG | 21.1 | 17.6 | -3.5 |
|  |  | 17.3 | -3.9 |
|  |  | 16.8 | -4.3 |
|  |  | 16.2 | -4.9 |

ND = difference in lag phase length was not detected

**Table S4.**
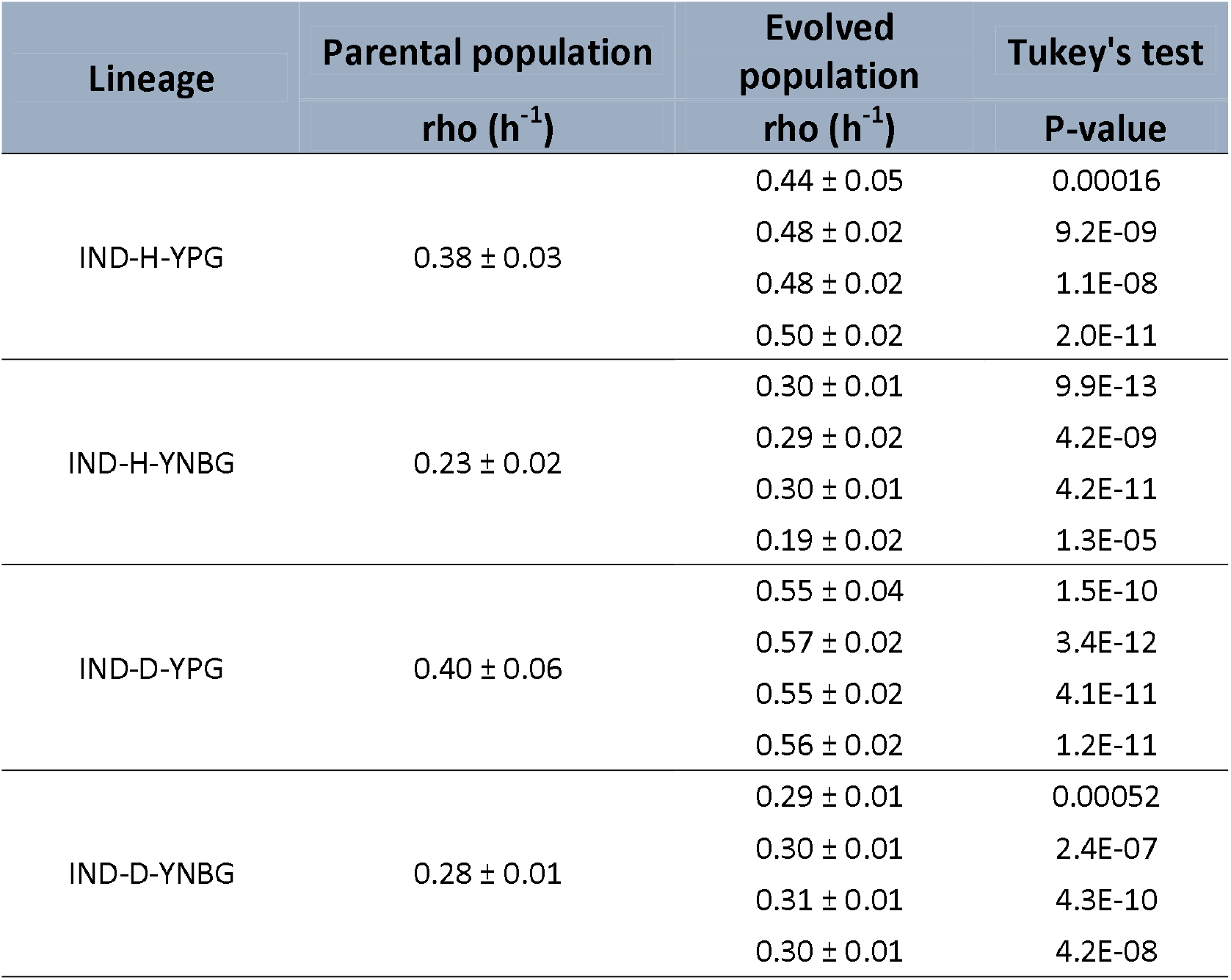
Maximum specific growth rate (rho) estimates of end-point populations of four replicate ALE lineages and respective parental populations of *S. cerevisiae* CEN.PK indigoidine pathway having (IND) haploid (H) and diploid (D) lineages adaptively evolved on YPG and YNBG media and grown in a multi-well plate format on the respective media. Tukey’s test P-values are shown for the differences between ALE end-point populations’ and the respective parental population’s maximum rho (one-way ANOVA and Tukey’s test (n_control_ = 10, n_case_ = 10), P-value < 0.01).

**Table S5.** Percentage of cell death in parental and adaptively evolved indigoidine synthesis pathway having (IND) *S. cerevisiae* CEN.PK haploid (H) and diploid (D) populations. The cell viability assessment was performed for cells grown on their respective adaptive evolution medium, i.e., either synthetic defined medium (YNBG) or rich medium (YPG) both with galactose as the carbon source. The assayed populations included parental and four adaptive evolution lineages after 25 transfers (25-1, 25-2, 25-3, 25-4). The data represents the means and standard deviations from triplicate measurements. P < 0.01 indicates a significant difference compared to the parental strain using ANOVA and Tukey’s test (n = 3).

| Lineage | Parental population<br>% of dead cells | Evolved population<br>% of dead cells | Tukey's test<br>p-value |
| --- | --- | --- | --- |
| IND-H-YPG | 9.6 ± 0.6 | 7.6 ± 0.1 | 5.6E-05 |
|  |  | 6.0 ± 0.2 | 3.1E-07 |
|  |  | 6.4 ± 0.2 | 9.2E-07 |
|  |  | 3.5 ± 0.2 | 1.0E-09 |
| IND-H-YNBG | 31.5 ± 0.2 | 17.6 ± 0.2 | 1.9E-11 |
|  |  | 15.9 ± 0.2 | 4.9E-12 |
|  |  | 24.1 ± 0.7 | 3.2E-09 |
|  |  | 26.8 ± 0.5 | 3.4E-07 |
| IND-D-YPG | 9.1 ± 0.3 | 8.2 ± 0.2 | 0.20 |
|  |  | 9.5 ± 1.0 | 0.88 |
|  |  | 4.8 ± 0.1 | 5.7E-06 |
|  |  | 8.3 ± 0.3 | 0.35 |
| IND-D-YNBG | 27.5 ± 0.4 | 45.2 ± 0.5 | 3.8E-11 |
|  |  | 25.1 ± 0.5 | 0.0018 |
|  |  | 23.5 ± 0.8 | 2.9E-05 |
|  |  | 24.2 ± 0.4 | 1.6E-04 |

**Table S6.** Maximum specific growth rate (rho) estimates of the end-point populations of four replicate ALE lineages and respective parental populations of *S. cerevisiae* S288C-based wild type (WT) and bikaverin pathway having (BIK) lineages adaptively evolved on glucose as carbon source on YPD and YNBD media and grown in a multi-well plate format in the respective media. Tukey’s test P-values are shown for the differences between ALE end-point populations’ and parental population’s maximum rho (one-way ANOVA and Tukey’s test (n_control_ = 10, n_case_ = 10), P-value < 0.01).

| Lineage | Parental population<br>$\rho$ ($\text{h}^{-1}$) | Evolved<br>population<br>$\rho$ ($\text{h}^{-1}$) | Tukey's test<br>P-value |
| --- | --- | --- | --- |
| WT-BIK-YPD | $0.17 \pm 0.01$ | $0.15 \pm 0.01$ | 1.4E-07 |
| | | $0.15 \pm 0.01$ | 3.1E-05 |
| | | $0.13 \pm 0.01$ | 7.8E-13 |
| | | $0.15 \pm 0.01$ | 2.3E-06 |
| WT-BIK-YNBD | $0.22 \pm 0.02$ | $0.22 \pm 0.02$ | 1.00 |
| | | $0.23 \pm 0.02$ | 1.00 |
| | | $0.20 \pm 0.01$ | 0.15 |
| | | $0.24 \pm 0.04$ | 0.55 |
| BIK-YPD | $0.21 \pm 0.05$ | $0.15 \pm 0.01$ | 2.5E-06 |
| | | $0.14 \pm 0.01$ | 6.1E-08 |
| | | $0.14 \pm 0.01$ | 5.2E-08 |
| | | $0.12 \pm 0.01$ | 1.3E-10 |
| BIK-YNBD | $0.21 \pm 0.02$ | $0.15 \pm 0.02$ | 0.012 |
| | | $0.16 \pm 0.02$ | 0.048 |
| | | $0.15 \pm 0.04$ | 0.025 |
| | | $0.21 \pm 0.07$ | 1.00 |

**Table S7.** Lag phase lengths of the end-point populations of four replicate ALE lineages and respective parental populations of *S. cerevisiae* S288C -based wild type (WT) and bikaverin pathway having (BIK) lineages adaptively evolved on glucose as carbon source on YPD and YNBD media and grown in a multi-well plate format in the respective media.

| Lineage | Parental population lag length (h) | Evolved population lag length (h) | $\Delta h$ |
| --- | --- | --- | --- |
| WT-BIK-YPD | 15.4 | 13.2 | -2.2 |
|  |  | 13.0 | -2.4 |
|  |  | 13.0 | -2.4 |
|  |  | 13.3 | -2.1 |
| WT-BIK-YNBD | 21.3 | 21.3 | 0.0 |
|  |  | 21.6 | +0.2 |
|  |  | 19.7 | -1.7 |
|  |  | 20.1 | -1.2 |
| BIK-YPD | 16.1 | 12.2 | -3.9 |
|  |  | 13.2 | -2.9 |
|  |  | 12.2 | -3.9 |
|  |  | 12.1 | -4.0 |
| BIK-YNBD | 26.2 | 22.9 | -3.3 |
|  |  | 23.1 | -3.1 |
|  |  | 23.5 | -2.7 |
|  |  | 22.3 | -3.8 |

**Table S8**. Variant calls in *S. cerevisiae* cen.pk based lineages with indigoidine pathway. Table_S8.xlsx

**Table S9.** Variant calls in *S. cerevisiae* S288C based lineages, wild type and with bikaverin pathway. Table_S9.xlsx

**Table S10.**
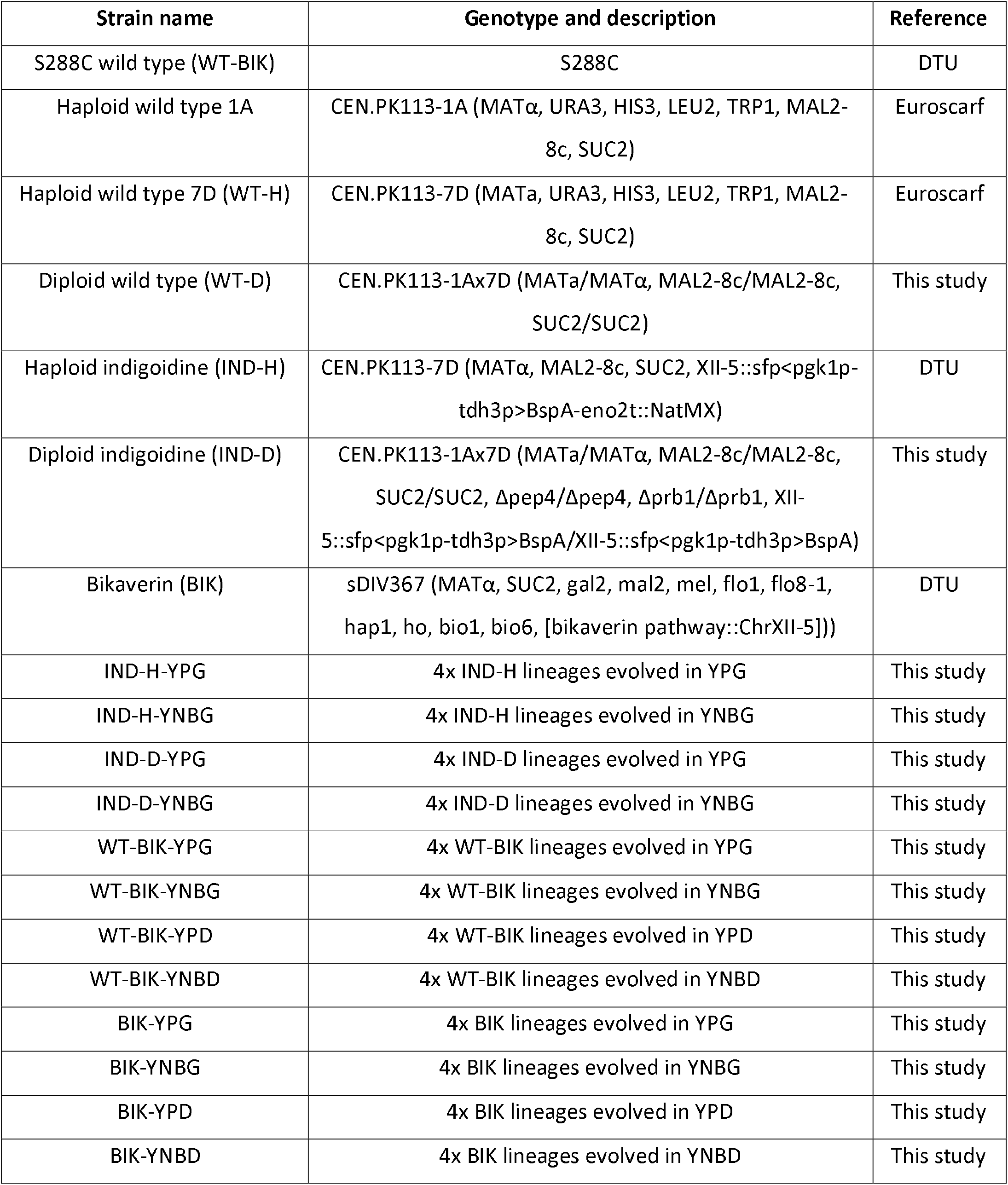
Strains used in this study.

**Table S11.** OD600 of yeast cultures used in cell viability analysis.

| Yeast cultures | OD600 |
| --- | --- |
| IND-D-YPG-0 | 36.2 |
| IND-D-YPG-25-1 | 35.2 |
| IND-D-YPG-25-2 | 34.5 |
| IND-D-YPG-25-3 | 35.7 |
| IND-D-YPG-25-4 | 30.5 |
| IND-D-YNBG-0 | 10.8 |
| IND-D-YNBG-25-1 | 16.3 |
| IND-D-YNBG-25-2 | 19.6 |
| IND-D-YNBG-25-3 | 18.1 |
| IND-D-YNBG-25-4 | 17.6 |
| IND-H-YPG-0 | 38.6 |
| IND-H-YPG-25-1 | 38.5 |
| IND-H-YPG-25-2 | 36.8 |
| IND-H-YPG-25-3 | 37.3 |
| IND-H-YPG-25-4 | 35.7 |
| IND-H-YNBG-0 | 13.8 |
| IND-H-YNBG-25-1 | 23.1 |
| IND-H-YNBG-25-2 | 23.2 |
| IND-H-YNBG-25-3 | 16.8 |
| IND-H-YNBG-25-4 | 17.2 |

**Table S12.** Whole genome sequencing sample accessions in ENA database.

| **ACCESSION** | **ALIAS** | **Sample title** |
| --- | --- | --- |
| ERS27024517 | BIK_YNBD_0 | BIK-YNBD-0 |
| ERS27024518 | BIK_ND_25_1 | BIK-YNBD-25-1 |
| ERS27024519 | BIK_ND_25_2 | BIK-YNBD-25-2 |
| ERS27024520 | BIK_ND_25_3 | BIK-YNBD-25-3 |
| ERS27024521 | BIK_ND_25_4 | BIK-YNBD-25-4 |
| ERS27024522 | BIK_YNBG_0 | BIK-YNBG-0 |
| ERS27024523 | BIK_N_25_1 | BIK-YNBG-25-1 |
| ERS27024524 | BIK_N_25_2 | BIK-YNBG-25-2 |
| ERS27024525 | BIK_N_25_4b | BIK-YNBG-25-4 |
| ERS27024526 | BIK_PD_25_1 | BIK-YPD-25-1 |
| ERS27024527 | BIK_PD_25_2 | BIK-YPD-25-2 |
| ERS27024528 | BIK_PD_25_3 | BIK-YPD-25-3 |
| ERS27024529 | BIK_PD_25_4 | BIK-YPD-25-4 |
| ERS27024530 | BIK_YPG_0b | BIK-YPG-0 |
| ERS27024531 | BIK_P_25_1b | BIK-YPG-25-1 |
| ERS27024532 | BIK_P_25_2b | BIK-YPG-25-2 |
| ERS27024533 | BIK_P_25_3b | BIK-YPG-25-3 |
| ERS27024534 | Y53_YNBD_0 | WT-BIK-YNBD-0 |
| ERS27024535 | Y53_ND_25_1 | WT-BIK-YNBD-25-1 |
| ERS27024536 | Y53_ND_25_3 | WT-BIK-YNBD-25-3 |
| ERS27024537 | Y53_ND_25_4 | WT-BIK-YNBD-25-4 |
| ERS27024538 | Y53_PD_25_1 | WT-BIK-YPD-25-1 |
| ERS27024539 | Y53_PD_25_2 | WT-BIK-YPD-25-2 |
| ERS27024540 | Y53_PD_25_3 | WT-BIK-YPD-25-3 |
| ERS27024541 | Y53_PD_25_4 | WT-BIK-YPD-25-4 |
| ERS27024542 | Y53_YPG_0 | WT-BIK-YPG-0 |
| ERS27024543 | Y53_P_25_3 | WT-BIK-YPG-25-3 |
| ERS27024544 | Y53_P_25_4 | WT-BIK-YPG-25-4 |
| ERS27024545 | IND_D_YNBG_0 | IND-D-YNBG-0 |
| ERS27024546 | IND_D_N_25_1 | IND-D-YNBG-25-1 |
| ERS27024547 | IND_D_N_25_2 | IND-D-YNBG-25-2 |
| ERS27024548 | IND_D_N_25_3 | IND-D-YNBG-25-3 |
| ERS27024549 | IND_D_N_25_4 | IND-D-YNBG-25-4 |
| ERS27024550 | IND_D_YPG_0 | IND-D-YPG-0 |
| ERS27024551 | IND_D_P_25_1 | IND-D-YPG-25-1 |
| ERS27024552 | IND_D_P_25_2 | IND-D-YPG-25-2 |
| ERS27024553 | IND_D_P_25_3 | IND-D-YPG-25-3 |
| ERS27024554 | IND_D_P_25_4 | IND-D-YPG-25-4 |
| ERS27024555 | IND_H_YNBG_0 | IND-H-YNBG-0 |
| ERS27024556 | IND_H_N_25_1 | IND-H-YNBG-25-1 |
| ERS27024557 | IND_H_N_25_2 | IND-H-YNBG-25-2 |
| ERS27024558 | IND_H_N_25_3 | IND-H-YNBG-25-3 |
| ERS27024559 | IND_H_N_25_4 | IND-H-YNBG-25-4 |
| ERS27024560 | IND_H_YPG_0 | IND-H-YPG-0 |
| ERS27024561 | IND_H_P_25_1 | IND-H-YPG-25-1 |
| ERS27024562 | IND_H_P_25_2 | IND-H-YPG-25-2 |
| ERS27024563 | IND_H_P_25_3 | IND-H-YPG-25-3 |
| ERS27024564 | IND_H_P_25_4 | IND-H-YPG-25-4 |

**Table S13.**
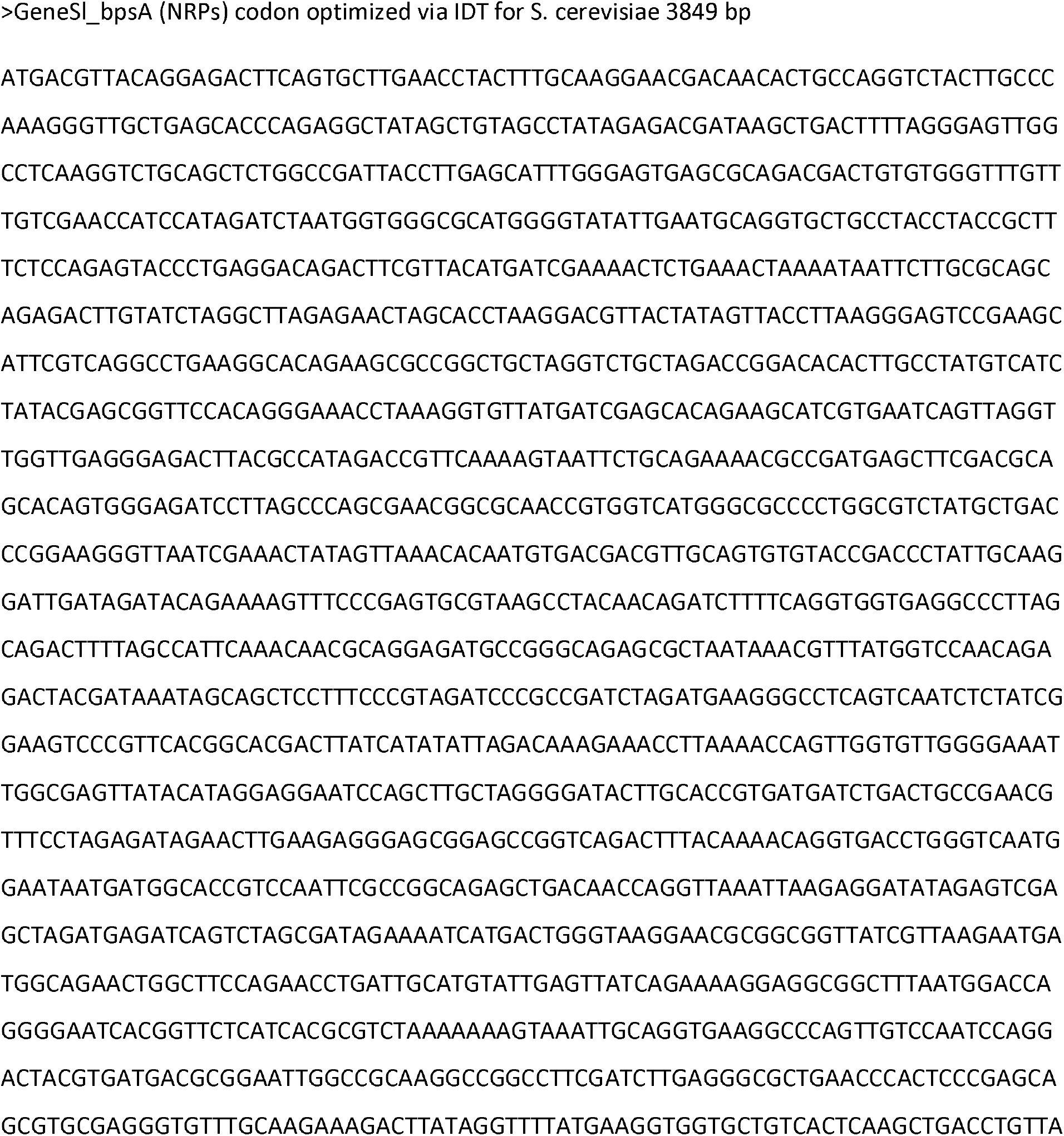
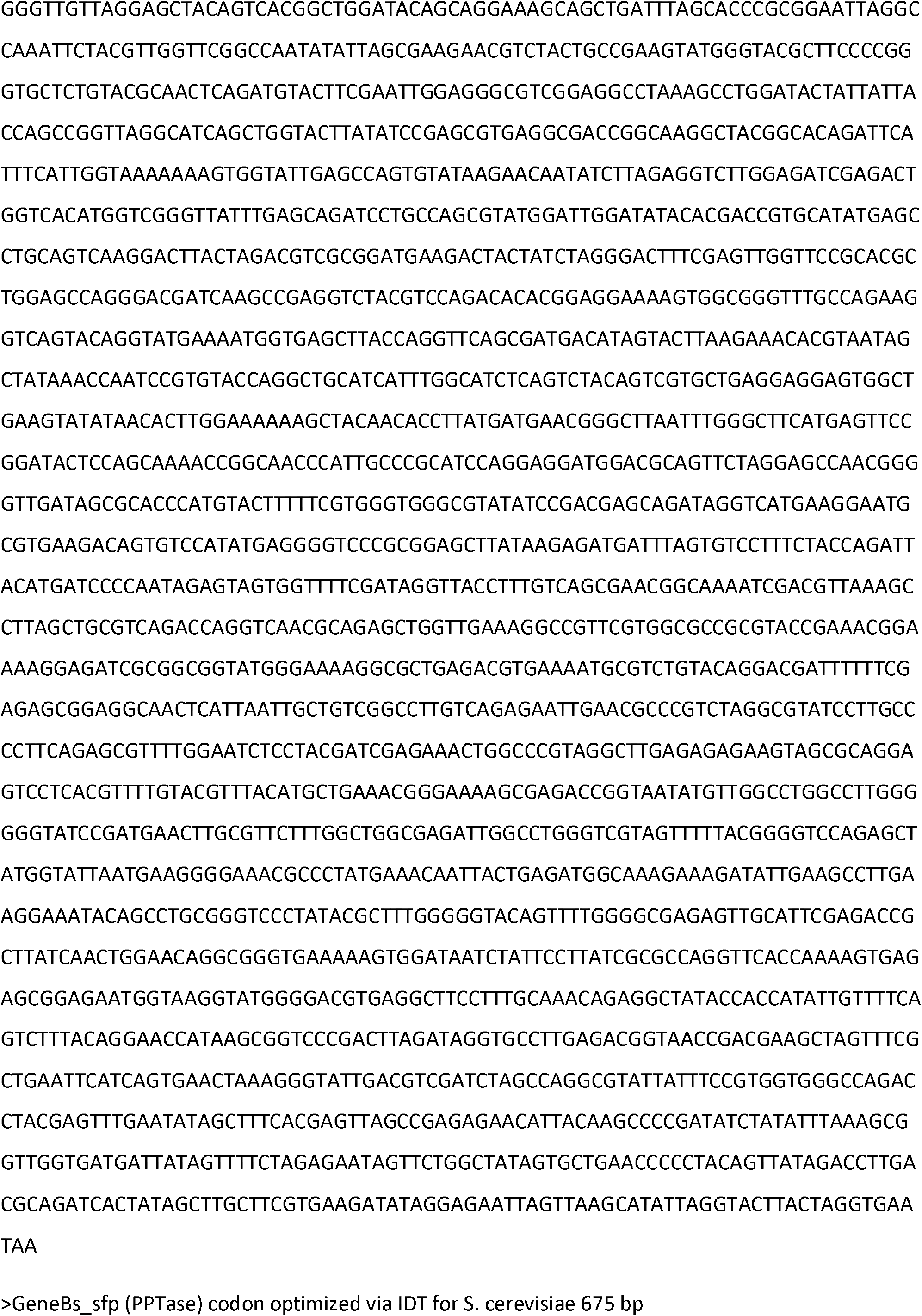
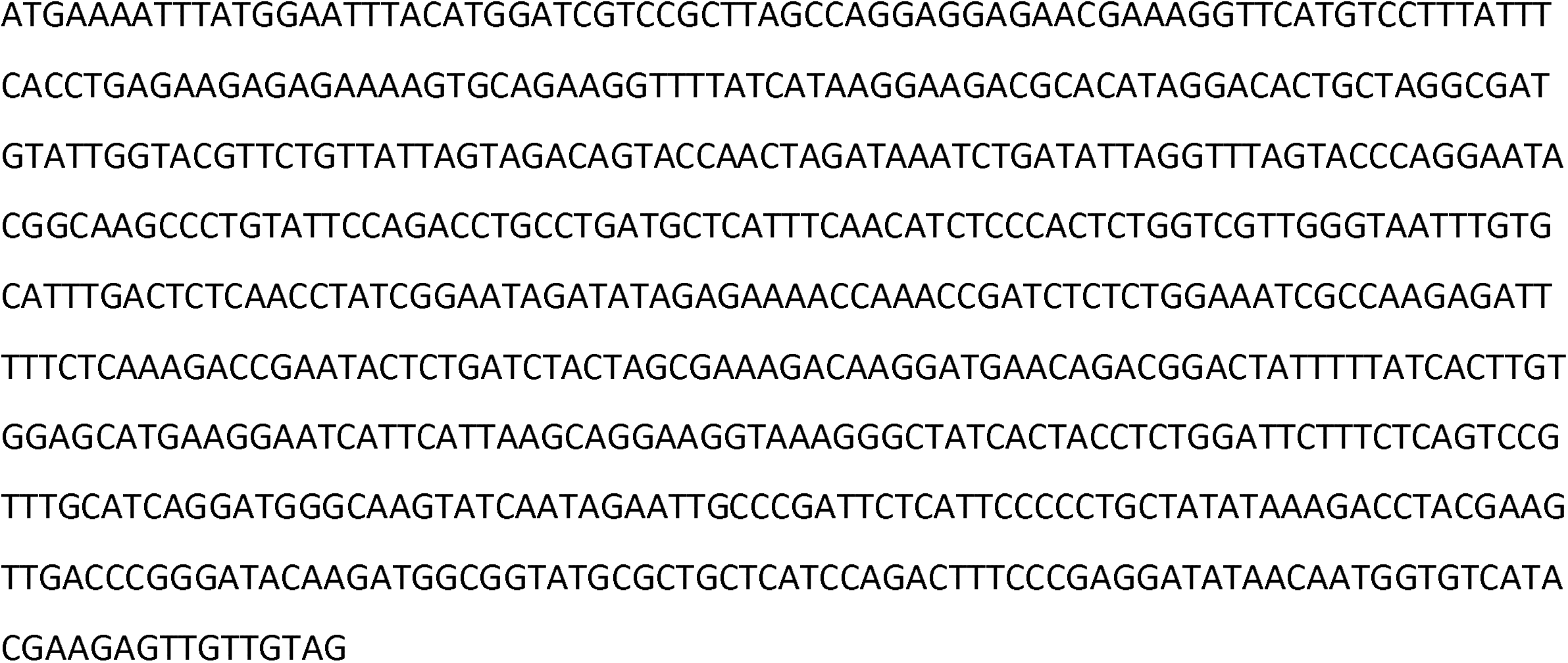
Reference sequences of indigoidine pathway genes.

**Figure S1.**
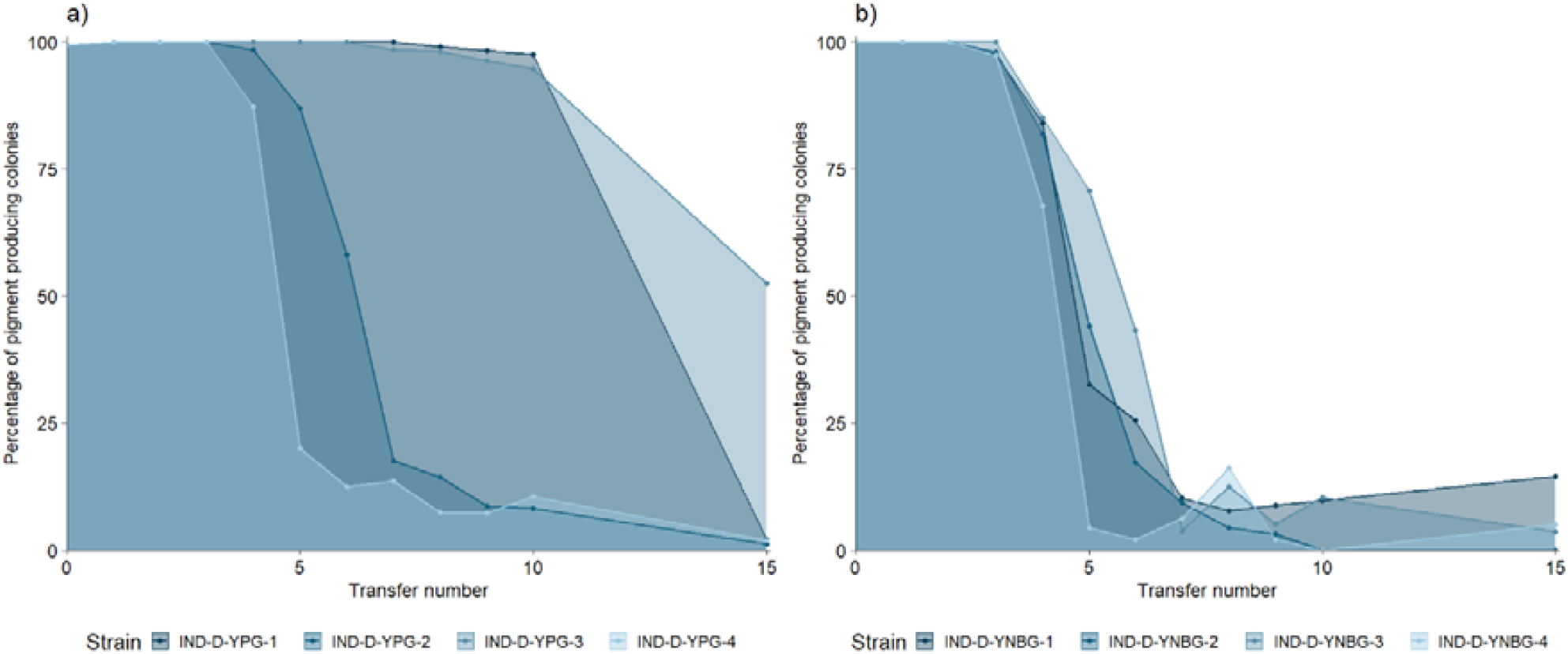
Percentages of indigoidine pigmented colonies in diploid *S. cerevisiae* CEN.PK lineages as a function of transfers in the ALE experiments in rich galactose medium (a) and in synthetic defined galactose medium (b) of having indigoidine pathway. Colony counting of all and pigmented colonies was performed for each ALE lineage on single plates except for the ALE end-point population in three replicates for each lineage.

**Figure S2.**
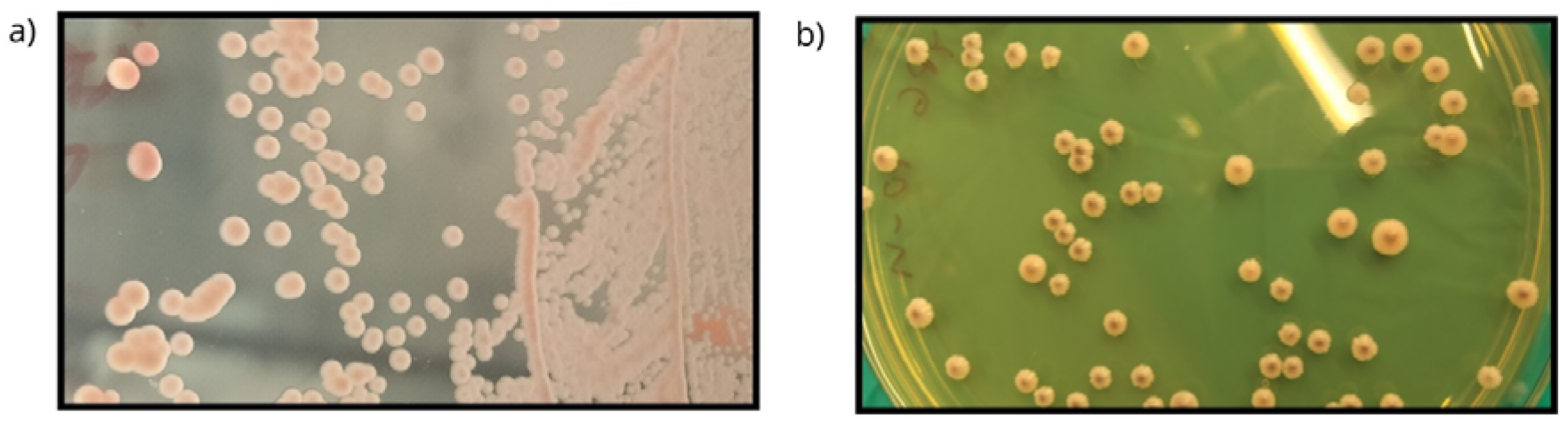
Colony pigmentation of haploid bikaverin pathway having *S. cerevisiae* S288C parental strain (a) and ALE lineage adaptively evolved for ten transfers in rich medium with galactose as the sole carbon source (b).

## Notes

### Competing Interest Statement

The authors have declared no competing interest.

