## Supplementary figures and images for "Adaptive evolution of engineered *Saccharomyces cerevisiae* in favored and unusual chemical environments"

### Supplementary material, Figure S1

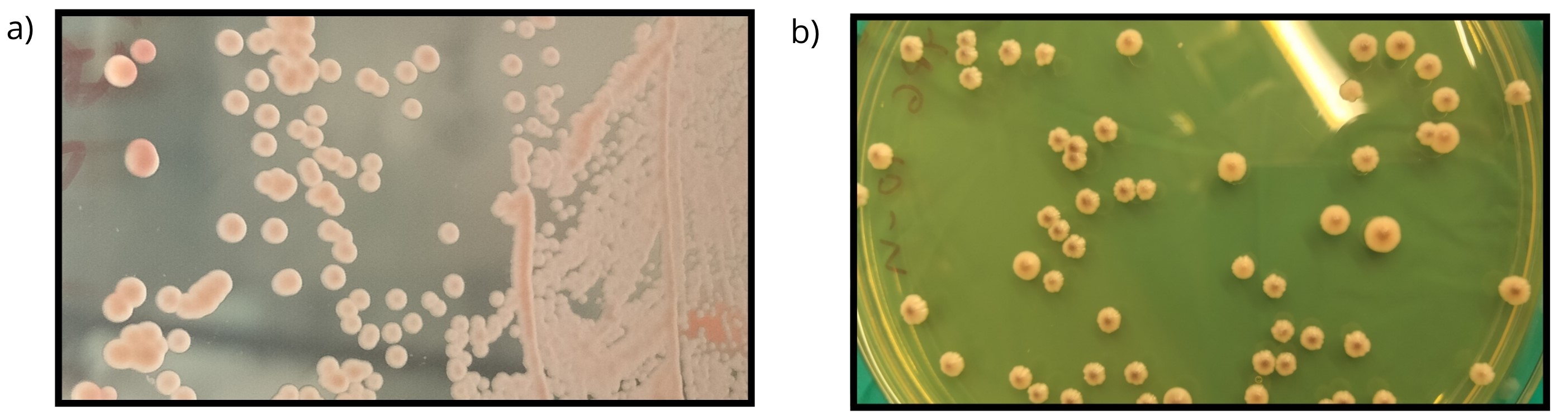

### Supplementary material, Figure S2

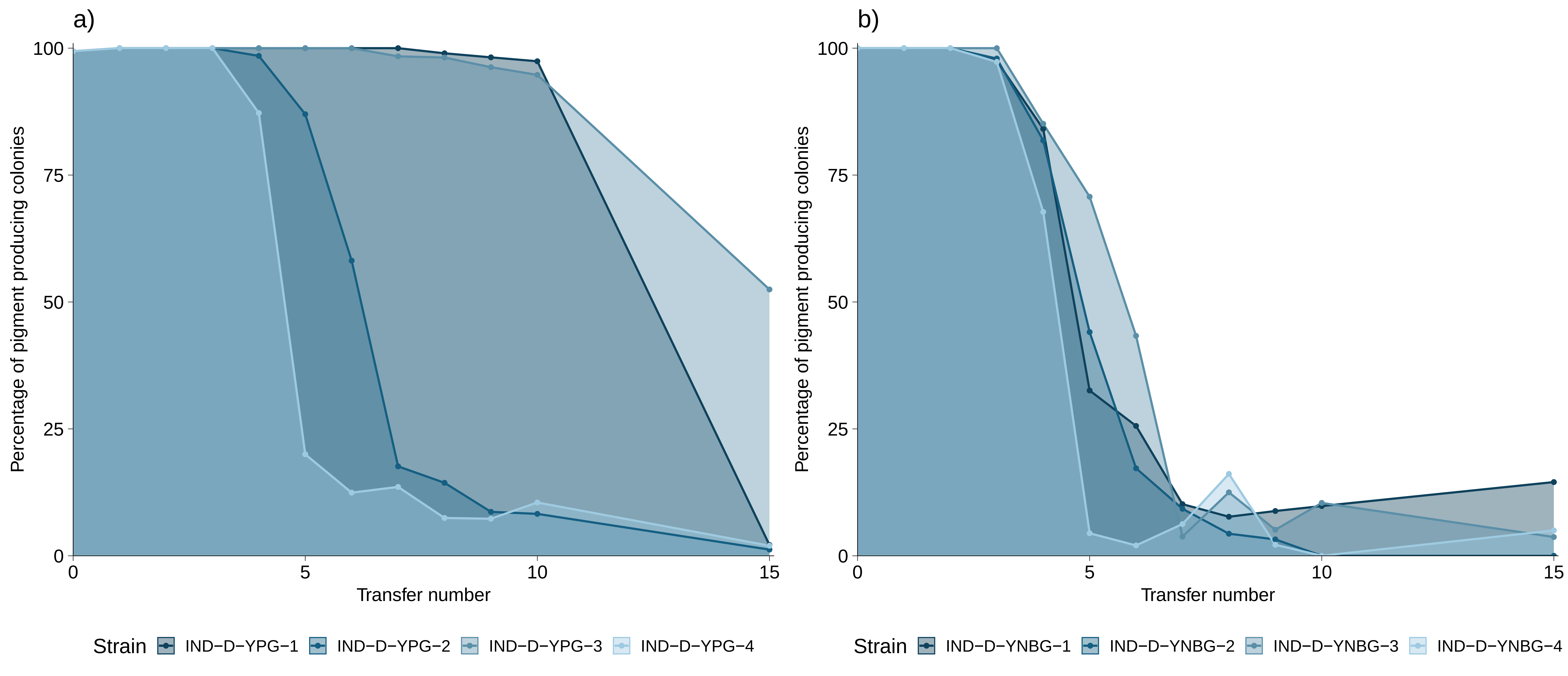
